# Chrono-atlas of cell-type specific daily gene expression rhythms in the regenerating colon

**DOI:** 10.1101/2025.11.20.689502

**Authors:** Vania Carmona-Alcocer, Cédric Gobet, Jessie MacDonald, Zainab Taleb, Felix Naef, Phillip Karpowicz

## Abstract

The circadian clock is a molecular timer present throughout the body, including the gastrointestinal tract, where it regulates daily rhythms in physiology through the timing of rhythmic gene expression. Dysfunctional rhythms, caused by loss of clock timing and/or environmental disruption is implicated with gastrointestinal dysfunction and pathology. The large intestine (colon) is composed of many different types of cells with distinct gene expression programs and functions. How daily rhythms in transcript abundance are coordinated in the intestine at a cell-specific level is not known. Using single cell transcriptomics, we analyzed 24-hour gene expression in all major cell types of the proximal and distal regions of the colon following injury. We find that daily gene expression is not uniform: rhythmic genes, including circadian clock components, clock targets, and systemic programs, differ in their timing and are cell-type specific. Cells of the epithelium, stroma, and immune system to display strong rhythms in metabolic, protein processing, and stress response genes during regeneration. While stromal and muscle cells exhibit robust circadian clock gene rhythms irrespective of injury, epithelial cells show weaker clock oscillations that become 12-hours antiphasic during regeneration. These data, completed with smFISH validation, reveal an unexpected complexity to daily transcript levels in the colon, and provide a resource for future studies by identifying the cellular source of 24-hour transcript rhythms.

## Introduction

The intestine is a complex organ composed of many different tissue cell types in very close proximity. These include cells of the epithelium, surrounding stroma, muscle, endothelium, lymphatics, enteric neurons and glia, and immune system^1–5^. Intestinal cells perform critical barrier functions and are among the most highly turned over cells in the entire body^6^. Notably, the epithelium is replaced continuously by a population of Intestinal Stem Cells (ISC) located at the base of the crypts of Lieberkühn^1^. Larger regions of tissue inflammation and injury are resolved by the efficient regeneration of the epithelial barrier that occurs through cellular dedifferentiation and tissue remodeling guided by the underlying stroma^7–11^. Such dynamic changes do not occur in a stable environment, rather the intestine is subject to significant variations arising from the daily consumption of nutrients and the subsequent metabolic responses by its resident microbiome^12–14^.

How cells coordinate their daily programs during the dynamic states of injury and regeneration is not known. Recent studies have implicated the circadian clock, a 24-hour molecular timer, as a major contributor to intestinal biology^15–20^. The clock drives circadian rhythms: 24-hour oscillations in physiology and behaviour^21^. The circadian clock is thought to be present in most cells and is thought to both synchronize them with the external daily environment, and to coordinate cell, tissue, and organ timing throughout the body. Circadian rhythms have been shown to impact human intestinal health^22–24^, and their disruption is implicated in Inflammatory Bowel Disease (IBD)^17,25–28^. Colitis, a form of IBD affecting the colon (large intestine), arises from episodes of acute inflammation that damage barrier cells which are then regenerated by ISCs to restore the tissue to its former state^11,29^. The loss of circadian rhythms has been tested in mouse models of Colitis that have shown clock function is required to resolve disease^30–34^. This implicates the circadian system in regulating intestinal physiological function and health.

Functionally the clock uses a near 24-hour transcription-translation feedback loop that drives rhythms in gene expression. It comprises the genes *Clock* and *Bmal1* whose products heterodimerize to drive the expression of their negative regulators *Per1-2* and *Cry1-2* ^21^. Twenty-four hour cycles of CLOCK/BMAL1 activity drive further components of the clock in the intestine, as well as thousands of genes throughout the body^35^. The intestinal microbiome also takes on circadian rhythmicity through food consumption: 24-hour feeding cycles cause the production of microbial metabolites that together with nutrient availability affect diurnal rhythms of gene expression in the host^13,36–38^. It is thought that these circadian and diurnal rhythms promote intestinal cell health by synchronizing the daily timing of gene expression with the needs of the body. Indeed, studies have revealed many 24-hour processes including digestion and absorption, cell proliferation, and inflammation have daily rhythms^13,36,37,39–43^. However, many questions remain. It is not clear whether all cells in the intestine possess circadian clock activity, or whether cells in this complex tissue exhibit synchronous gene expression rhythms relative to each other. Although diurnal rhythms are present, it is not clear which cells are responsible for these, or how or if these rhythms would be affected by the dynamic cell turnover occurring during regeneration. For instance, it has been proposed that undifferentiated Intestinal Stem Cells (ISCs) do not have clock activity but instead respond to circadian rhythms from the nearby differentiated Paneth cells, whose clock function drives rhythmic signal transduction in ISCs^44^. On the other hand, stem cells in other tissues have been shown to exhibit robust circadian clock function^45,46^. Elucidating the cell-specific rhythmicity in this tissue, and whether it is altered during regeneration, is key to understanding how daily intestinal physiology is regulated.

To address gene expression rhythms at the single cell level we sequenced the regenerating colon following injury to generate a chrono-atlas of thousands of rhythmic mRNA profiles for >20 different cell types of this organ. Our results reveal that robust circadian clock gene rhythms are strongest in the cells of the stroma, muscle, and endothelium which exhibit synchronous timing. However, other cell types exhibit varying degrees of circadian rhythmicity. This is most evident in the epithelial cells which display weak circadian clock oscillations in undamaged tissue that become antiphasic (12-hour opposite) relative to the surrounding cells during regeneration. We discover that diurnal oscillation of transcripts includes genes involved in metabolism, thermal-responses, and protein production / recycling, and assign specific signaling and cellular pathways to all major cell types in proximal and distal regions of the colon. Regeneration is accompanied by biphasic 24-hour gene rhythms, that peak in the morning and evening, suggesting a reprogramming following injury that renders the tissue susceptible to daily systemic physiological rhythms. Our data provide insight into the specificity of transcriptional rhythms in a cell-specific manner, highlighting the complex organization of daily cellular timing occurring within cells in this rapidly regenerating and environmentally responsive tissue.

## Results

### Circadian rhythms are present during colon regeneration

The mouse colon consists of a proximal (in humans ascending) region and distal (in humans descending) region, hence, we first tested the effects of the chemical Dextran Sulfate Sodium (DSS) in inducing a proximal *vs.* distal colon injury / regenerative response^29,47,48^. DSS was applied for 7 days to the *Per2-Luc* mouse strain, that contains a Luciferase enzyme fused to the *Per2* clock gene^49^, then withdrawn. Mice were tested at different stages pre- and post-DSS exposure to monitor damage and subsequent regeneration (Supplementary Fig 1). The distal colon region shows normal undamaged morphology at day 0, prior to DSS-exposure, which progresses to lesioned (ulcerated and crypt-absent) morphology at days 7-10, and large amorphous crypts indicative of tissue remodelling at days 10-17 (Fig 1A-B, Supplementary Fig 1B-C). Of note, the proximal colon is not appreciably affected and does not show the damage or regeneration present in the distal colon at the DSS levels provided in these tests (Fig 1A-B, Supplementary Fig 1B-C). Staining for Ki67, to test regenerative proliferation, confirmed that regeneration is induced in the distal but not proximal colon at days 10-17 (Fig 1C-D). Lesions in the distal colon are highest at day 10, 3 days upon DSS withdrawal, and begin to resolve as regeneration increases at day 17 as the organ recovers from injury. These data are consistent with other reports studying the resolution of colitis through tissue regeneration following DSS administration^29,47,48^. We therefore used the distal region as a system to study daily rhythms during injury / regeneration, and the proximal region as an internal control of undamaged colon tissue.

**Figure 1.**
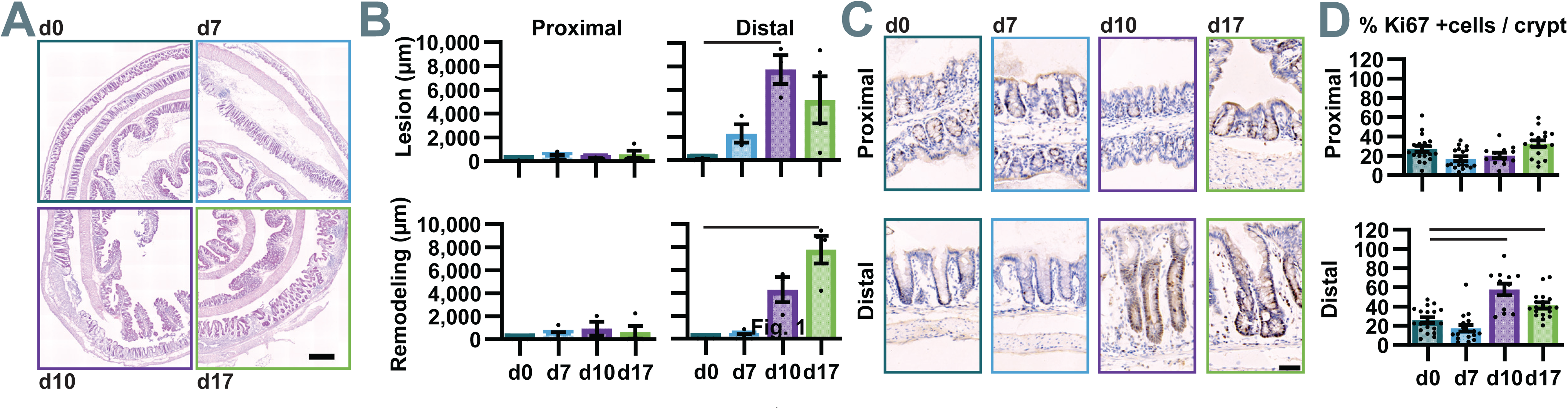
Temporal dynamics of colonic injury and regeneration. **A)** Representative Hematoxylin and Eosin-stained images of colon tissue at day 0 (no damage), day 7 (end of DSS treatment), day 10, and day 17 (recovery phase). Scale bar = 500 µm. **B)** Quantification of total lesions (upper graphs) and regenerating regions (lower graphs) indicates that DSS induce colitis mainly affects the distal colon, with greatest severity at day 10 and increased regeneration at day 17. **C)** Representative images of anti-Ki67 immunostaining (brown) in crypts from the proximal (upper images) and distal (lower images) colon. Scale bar = 50 µm. **D)** Percentage of proliferating Ki67+ cells is stable in the proximal colon but increases in the distal colon at days 10 and 17. Statistical comparisons were performed using Kruskal-Wallis test with Dunn’s post hoc test relative to day 0. Data are presented as mean ± s.e.m. (n=3 mice by timepoint). Significant differences (p < 0.05) are indicated by black bars.

We asked whether daily behavioural rhythmicity is lost during injury and/or during the regenerative response. To test this, *Per2-Luc* mice were treated as before and maintained in behavioural chambers to continuously monitor circadian locomotor activity and feeding, two measures of an individual’s daily rhythms which are regulated by the circadian clock (Supplementary Fig 2A). In mice kept under a 12-hour light / 12-hour dark photoperiod with *ad libitum* access to food, locomotor activity follows a diurnal rhythm pre- and post-DSS exposure, with a slight reduction in the amplitude of behavioural activity rhythms occurring during the acute phases of injury at days 4-10 (Fig 2A-D, Supplementary Fig 2B-E). Feeding follows a strong diurnal rhythm with no change in amplitude (Fig 2E-I), even though a reduction in the total daily food consumed occurs at days 5-10 (Fig 2F). These data indicate that daily activity and feeding rhythms are intact during injury and regeneration, even persisting during the heightened stages of illness present due to DSS administration.

**Figure 2.**
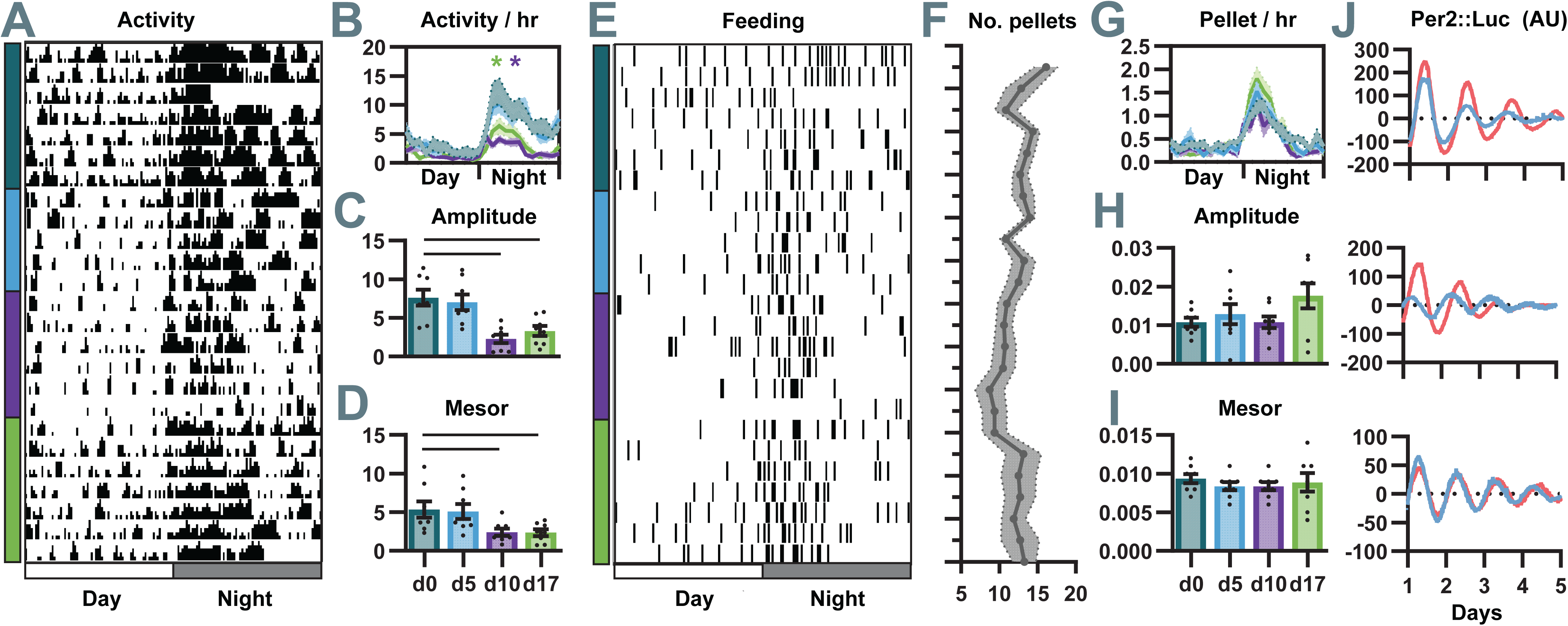
Circadian rhythms are present during colon regeneration. **A)** Representative single-plot actogram of locomotor activity of one individual mouse (each line indicates one 24 hour day). Behavior demonstrates 24-hour patterns of rest and active times. Color code indicates analysis sections corresponding to disease stages: dark blue = no damage, light blue = no disease symptoms, purple = disease symptoms apparent; green = recovery phase. **B)** Mean daily locomotor activity profiles indicates mice are active at night, but show reduced activity during injury and recovery. Analysis reveals a significant decrease in amplitude **C)** and mesor **D)** during injury and recovery. **E)** Representative single-plot actogram of food intake rhythms for one individual mouse (each line indicates one 24 hour day). **F)** Mean daily pellet intake remains consistent across disease stages with a slight decrease in total amount ingested during injury phases. **G)** Mean daily food intake profiles across disease stages shows most food is consumed at night during peak activity. Rhythmic analysis of food intake shows no significant changes in either amplitude **H)** or mesor **I)** indicating 24-hour daily feeding rhythms a preserved over injury and recovery phase. Behavioral data were collected from n=8 mice. Data are presented as mean ± s.e.m. **J)** *Per2::Luc* bioluminescence subtracted traces from distal proximal (blue) and distal (red) colon explants collected at ZT01 on day 17 indicate circadian oscillations persist in the regenerating colon (n=3 representative mice shown). Daily profiles (b, e) were analyzed using repeated-measurements two-way ANOVA, comparing each curve to a baseline curve (undamaged). Circadian parameters (c, d, h, i) were analyzed by one-way ANOVA with Dunnett’s post hoc test relative to day 0. Significant differences (p < 0.05) are indicated with asterisks or black bars.

Our previous data has shown rhythms are diminished in the colon tissue during acute injury^30^. To further confirm that regenerating colon tissue itself exhibits circadian clock function 10 days post-injury, we tested colon tissues of *Per2-Luc* mice at day 17 following DSS administration. At this stage both lesions and remodelling tissue are evident in all tissue samples collected over 24 hours in the distal but not proximal colon (Supplementary Fig 2E). Both distal and proximal regions display near 24-hour free running circadian clock rhythmicity when explants are monitored for luminescence over 6 days in culture (Fig 2J). We conclude that circadian clock rhythms are present in regenerating colon tissue, just as behaviour and feeding rhythms are present at the individual level.

### The regenerating colon exhibits biphasic mRNA rhythmicity

Given the persistence of circadian rhythms during regeneration, we asked what trancripts show daily rhythmicity in their levels in the different cells of this complex organ. We investigated cell-type-specific diurnal rhythms in the regenerating colon using single-cell RNA-seq (scRNA-seq). Proximal and distal colon were harvested from mice at day 17 post-injury over four Zeitgeber Times (ZT, refers to time with respect to photoperiod with ZT0 = lights turn on, ZT12 = lights turn off). Tissues were flash-frozen upon collection at ZT01 / ZT07 / ZT13 / ZT19 and subsequently processed and sequenced using the 10x Genomics Chromium platform with the Fixed RNA profiling assay (Flex-seq, Fig 3A). After filtering, we obtained on average 5,500 cells per sample with a median of 2,100 UMIs per cell. Cell clustering was performed to identify tissue cells present in the colon including epithelial, immune system, smooth muscle, stromal, endothelial, lymphatic, and enteric nervous system (Fig 3B-C). These cells were subdivided further into 30 cell-specific clusters which were identified based on previously published cellular markers (Fig 3D, Supplementary Fig 3A-D)^29,50–62^. All major classes of colon epithelial cells were detected, from Intestinal Stem Cells (ISC) to their differentiated progeny. ISCs were subdivided into a separate cluster of active-proliferating ISC subtypes (ISC-proliferative) *vs*. less proliferative^63^, and both Goblet and Colonocyte precursors (Transit Amplifying, TA) were discriminated from Goblet cell and Colonocytes, respectively (Fig 3D). Enteroendocrine and Tuft cells were also detected. The supporting Stromal tissues that underlie the epithelium were found and subdivided into Reticular cells, Interstitial cells, Trophocytes, and Telocytes. Similarly, muscle was subdivided into Muscularis Mucosa (MM), Muscularis Propria (MP), and Lamina Propria Myocytes (LPM) (Supplementary Fig. 4A-D). Cells of the Immune System, that are numerous in the colon, were identified including Macrophages, Dendritic cells, T-cells and actively dividing T-cells (T-proliferative), B-cells and dividing B-cells (B-proliferative), and Plasma cells. Endothelial cells, Lymphatic cells, and both Enteric Glia and Enteric Neurons called the Interstitial Cells of Cajal (ICC) were also identified. Although there was some variability in the total number of cells based on the time of sample collection (Fig 3E), and their relative abundance in proximal *vs*. distal regions (Fig 3E-F), cell-types were evenly proportioned across ZT01 / ZT07 / ZT13 / ZT19 experimental samples in all clusters. The representation of cell types across samples supports the use of these data to detect 24-hour time-dependent changes throughout the colon.

**Figure 3.**
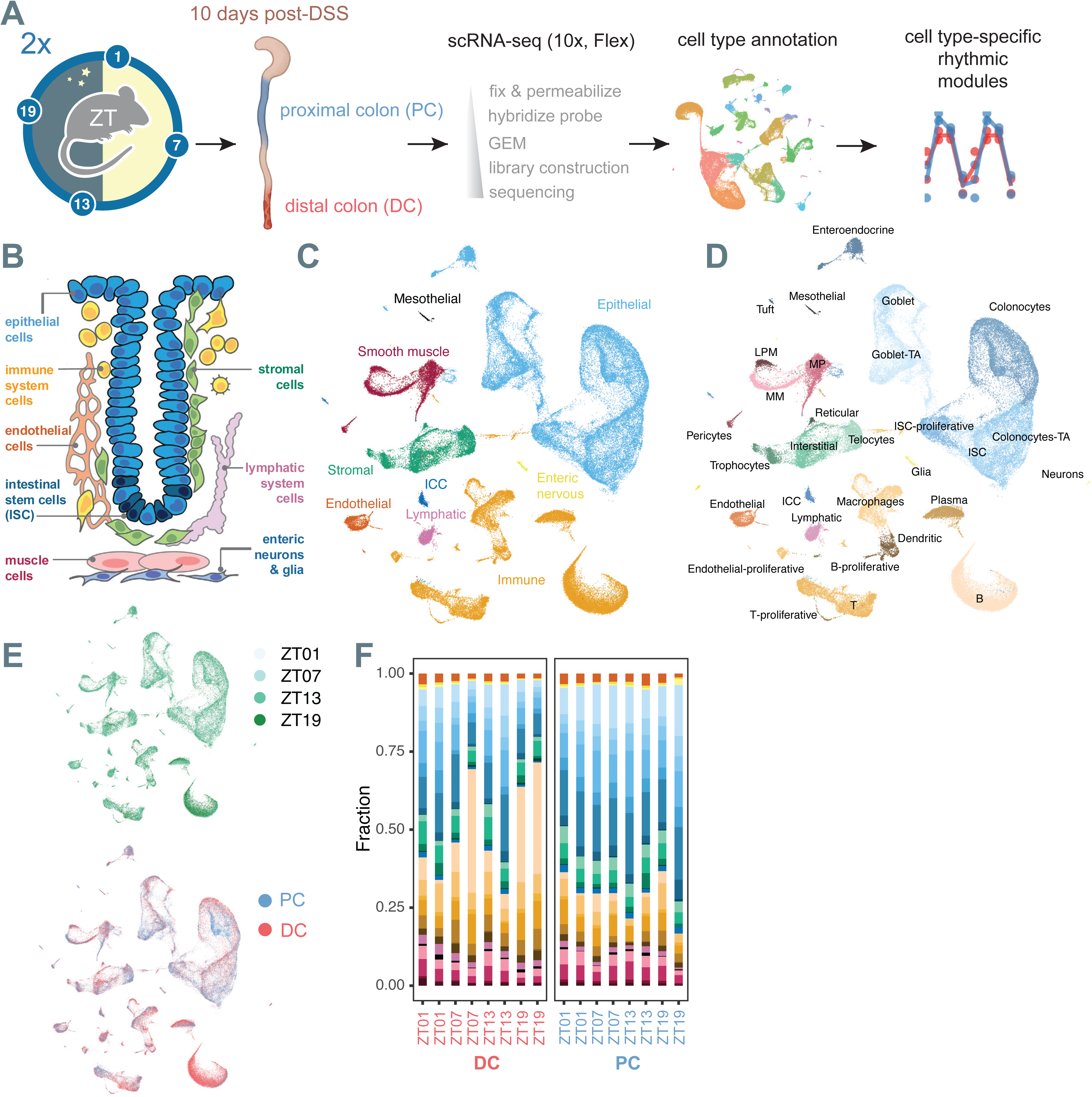
Twenty-four-hour scRNA-seq identifies distinct colon cell types. **A)** Schematic of experimental design. Mice were treated with DSS and sacrificed at 10 days post-treatment at four time points (ZT as indicated). Proximal colon (PC, blue) and distal colon (DC, ref) tissues were collected and processed for scRNA-seq using 10x Genomics Flex, followed by cell type annotation and cell type-specific 24-hour rhythmic assessment. **B)** Schematic depicting the major cell types identified in the scRNA-seq with their respective color-coding. **C)** UMAP visualization of all integrated cells colored as in panel B. **D)** Detailed UMAP of all cell-specific subsets showing 30 subpopulations detected. E) UMAP color-coded by time (ZT01, ZT07, ZT13, ZT19) and proximal (PC) or distal regions (DC). **F)** Fraction of cells detected by colonic region in the samples / regions tested showing the relative proportions of each cell type in distal (DC) versus proximal colon (PC) across time points. Overall, cell types were represented across regions and timepoints, although some variability in the numbers of certain cells such as B-cells was noted.

Next, 24-hour rhythmicity of gene expression was assessed in cell types with sufficient representation across samples. Of the 30 cell types identified, 21 with a minimum of 1000 cells were included in this analysis. Using pseudo-bulk mRNA levels in each sample to control for biological variability, we then analyzed rhythmicity in these cell types in both the proximal and distal colon, applying harmonic regression to log-normalized counts. To select for both significantly rhythmic patterns and rhythms of sufficient amplitude, genes having a log_2_ amplitude > 0.5 and p-value <0.05 (n=8 pseudo-bulk samples) were considered for further analysis (Supplementary Table 1). With these criteria, a total of 9,539 unique genes exhibited 24-hour rhythmicity in at least one cell type or region, with the distal regenerating colon exhibiting more compared to the proximal colon (Fig 4A, Supplementary Fig. 5A). More rhythmic genes are present in the epithelial lineages than in any other types of cells, nonetheless, rhythmic genes were detected in most of the cell types found (Fig 4A). In previous studies, diurnally rhythmic genes can peak in the colon at various times over a 24-hour period^13,37,39^, hence we examined whether the timing of cell-type specific rhythms are equally distributed over the day. In the regenerating colon the phase distribution of rhythmic genes is spread over 24-hours, with genes peaking at different times throughout the day in endothelial cells, and ICCs (Fig 4B). However, other cells such as the epithelia, stroma, immune system, lymphatic, and muscle exhibit biphasic rhythms with most genes peaking at either the morning (ZT0-4) or evening (ZT12-16) (Fig 4B). These times correspond to when mice begin their sleep (morning) or wake (evening) and begin to feed (Fig 2). In the proximal undamaged colon, this pattern is different: endothelial cells and ICC neurons exhibit peak gene expression in the midday (ZT4-8), and this same pattern is also evident in epithelia, stroma, immune system, lymphatic, and muscle (Fig 4C). This midday timing occurs during periods when nocturnal mice are inactive and fasting. The state of regeneration thus appears to correlate with a transcriptional reprogramming in daily gene expression timing. Indeed, it is clear in epithelial and stromal cells that show distinct gene rhythm timing in regenerating (distal) *vs*. non-regenerating (proximal) conditions (compare Fig 4B-C). These results suggest that distal regenerating cells exhibit a biphasic gene expression pattern that correlates with behavioural activity and feeding *vs.* rest.

**Figure 4.**
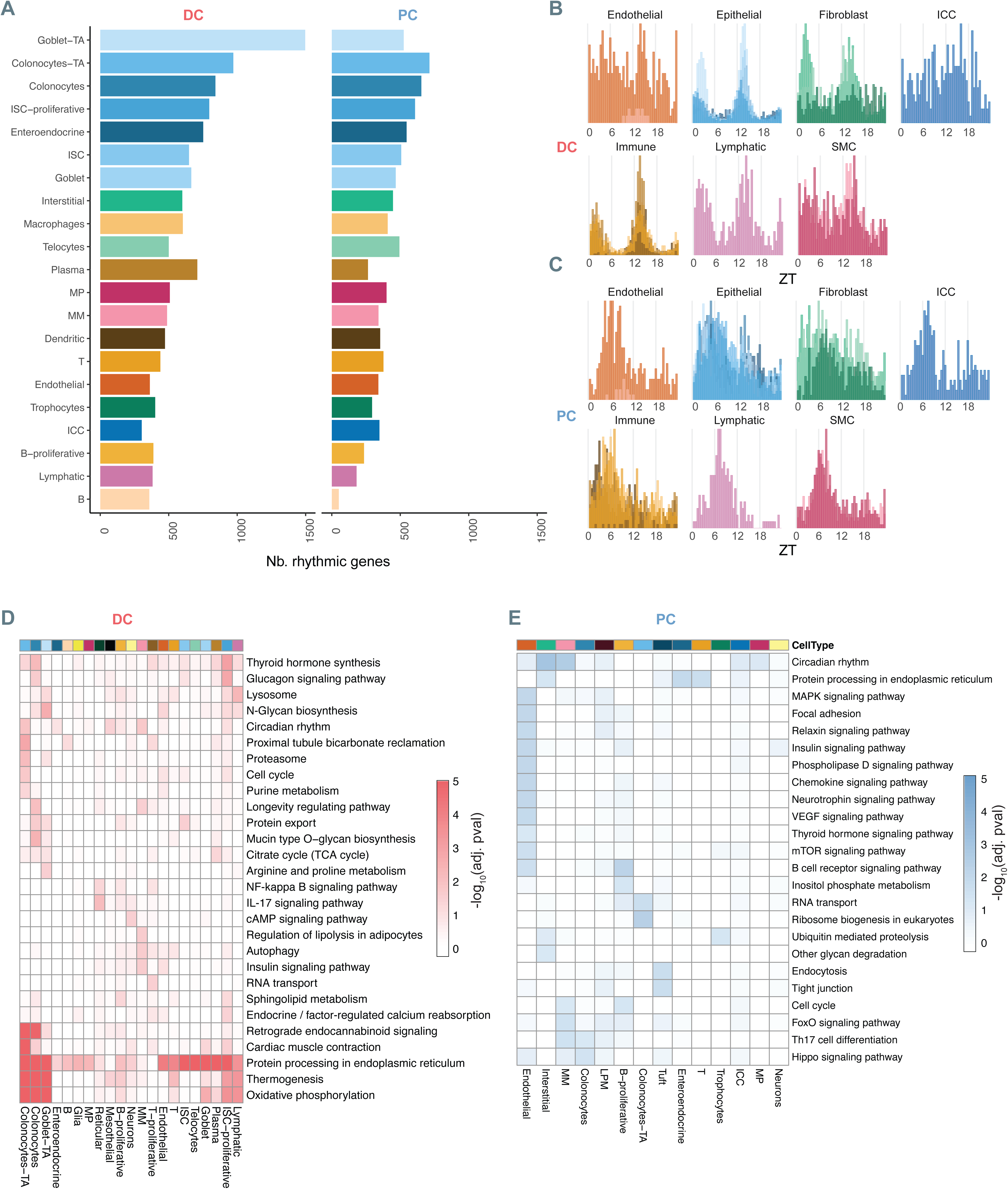
The regenerating colon exhibits mRNA rhythmicity and reprogramming in the regenerating colon. **A)** Number of rhythmic genes in each cell type across the distal (DC) and proximal (PC) colon. Rhythmicity was determined by harmonic regression (p-value < 0.05; log₂ peak-to-trough amplitude > 0.5). **B-C)** Phase distribution of rhythmic genes according to time of peak expression in each cell type in the distal and proximal colon. **D)** Heatmap showing significantly enriched KEGG pathways in rhythmic genes across cell types in the distal colon. Colors represent -log₁₀(adjusted p-value). **E)** Same as in (D), for the proximal colon.

### Regeneration reprograms daily rhythms to metabolic and cell stress responses

To further investigate these changes, we first compared rhythmic gene expression programs between regenerating and non-regenerating colon across all cell types. Rhythmic genes were profiled by pathway enrichment (KEGG) to detect 24-hour cellular changes occurring in 21 cell types under these conditions. The pathways in the regenerating distal colon show little overlap with the non-regenerating proximal colon, suggesting that different programmes are elicited (Fig 4D-E). The regenerating distal colon shows rhythms in many metabolic processes including oxidative phosphorylation and the TCA cycle, thermogenesis, lipolysis, arginine/proline, purine, and sphingolipid metabolism (Fig 4D). Enriched insulin, glucagon, cAMP, and thyroid signaling pathway rhythms indicate tissue responses that may be subject to daily feeding. In addition, rhythmic cellular processes such as autophagy and proteasome function, and the enrichment of NFκB and IL-17 signaling suggest a 24-hour stress and/or inflammation response. Endoplasmic reticulum protein processing, and N-glycan and mucin biosynthesis are rhythmic in many epithelial cells – highlighting different aspects of protein production and intestinal mucous barrier function that also exhibit daily rhythms.

In contrast to the regenerating colon, the proximal undamaged colon has a different rhythmic gene programme enriched in many cellular signaling pathways. 24-hour rhythms in MAPK, VEGF, Neurotrophin, FoxO, and Hippo pathways in the undamaged colon suggest that cellular communication follows a daily cycle (Fig 4E). Cellular processes under 24-hour rhythms included Phospholipase D and Inositol pathways, RNA transport and ribosome biogenesis, Endocytosis, and Tight junctions. B cell receptor and Chemokine signaling, and Th17 cell differentiation were also detected, indicating daily immune system regulation. Like the regenerating colon, pathways associated with feeding are also represented including mTOR, Insulin, and Thyroid signaling. Consistent with previous studies, the colon has significant 24-hour rhythms in transcript levels in many cellular and signaling processes^13,37,39^.

Altogether, these data suggest that regeneration alters daily rhythms in cellular signaling pathways to drive rhythms in metabolism and cellular stress responses. However, some pathways are shared between regenerating and undamaged regions, for instance circadian rhythm genes are detected in both indicating that clock function is present at the overall tissue level.

### Cell-type specific mRNA rhythms are linked with diverse functions in the 24-hour colon

We further probed subtypes of cells, such as the ISCs and stroma that underlie regeneration, to categorize transcripts detected in the proximal and distal colon regions. Pathway analysis (KEGG, REACTOME, Wiki Pathways) indicates that ISCs are enriched for rhythmic genes involved in Notch signaling, fatty acid metabolism, and cell cycle regulation (Fig 5A). The more actively proliferative ISCs (Fig 3D) are enriched for mitochondrial genes involved in oxidative phosphorylation and thermogenesis, with genes such as *Atp5e* and *Cox6a1* showing stronger rhythms in the regenerating distal colon than the undamaged proximal colon (Fig 5A-B). Of note, transit amplifying precursors (Colonocyte TA, Goblet TA) and differentiated colonocytes and goblet cells share this enrichment for mitochondrial components (Fig 5C), suggesting daily rhythms in aerobic oxidation are common to epithelial cell types. On the other hand, Tuft cells are enriched for genes involved in endocytosis, tight junctions, and Egf / MapK signaling (Supplementary Fig 6A) indicating these types of epithelial cells exhibit unique cell-specific cellular timing. Overall, the epithelial cell lineages exhibit both shared and unique 24-hour rhythms in their gene expression profiles.

**Figure 5.**
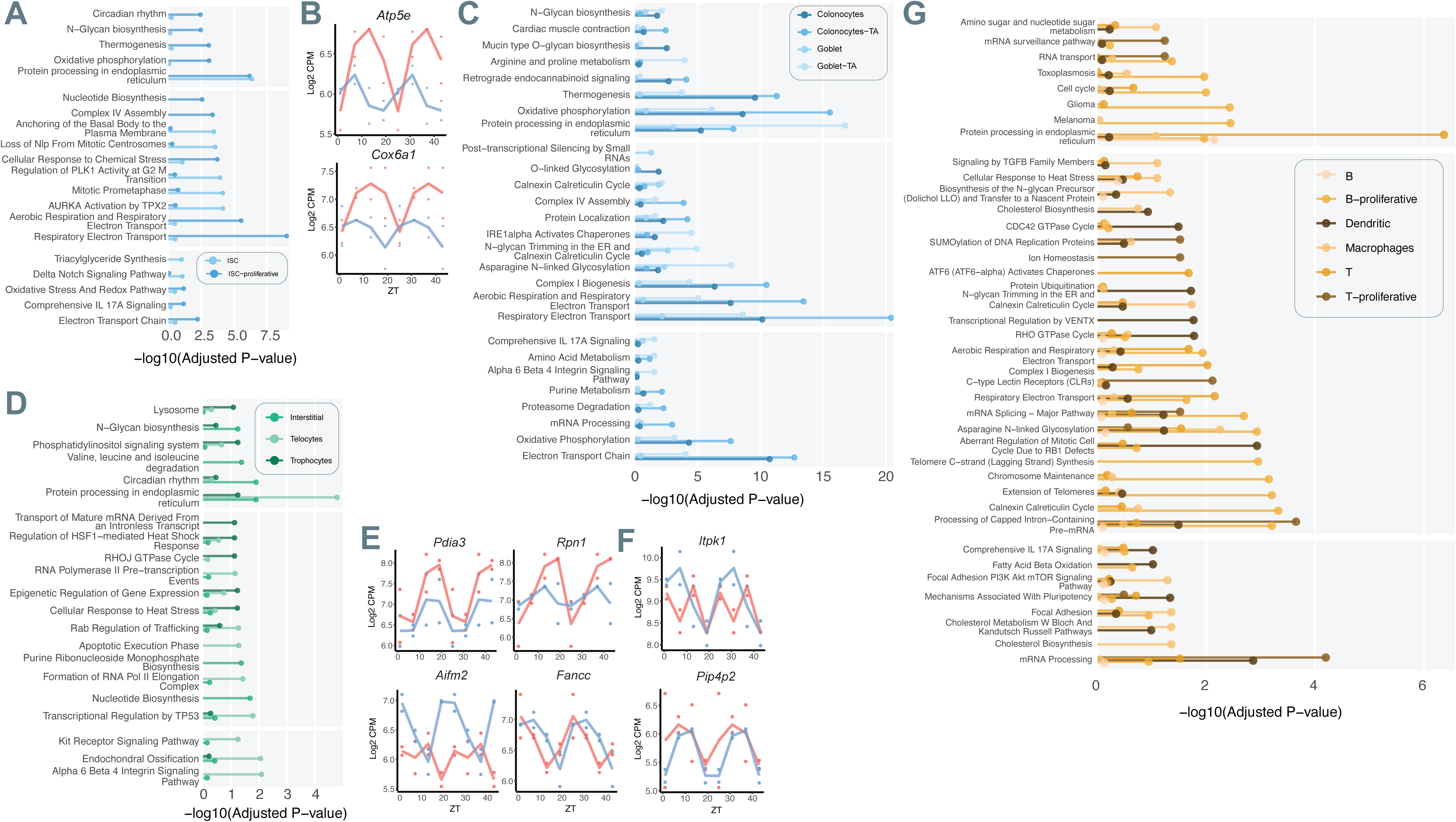
Cell-type specific rhythms in gene expression reveal diverse transcriptional programs in the 24-hour colon. Enrichment of rhythmic genes in specific cell types for major biological pathways. Bar plots show the negative log_10_ of the adjusted P-value (-log_10_ adjusted p-value) from Enrichr analysis (KEGG, REACTOME, WikiPathways) for genes exhibiting 24-hour rhythms (p-value<0.05 and Log 2 peak-to-trough amplitude >0.5) in each specified cell type. **A)** Enrichment in ISC and Proliferative ISC (ISC-proliferative). **B)** *Cox6a1* and *Atp5e* normalized expression level (Log_2_ CPM) in ISCs shows strong 24-hour rhythms particularly in the regenerative distal colon (DC, red) compared to the proximal colon (PC, blue). Dots represent the pseudo-bulk normalized count for each individual animal at a given ZT point, and the colored line shows the average expression trend for the Proximal Colon (PC, blue) and Distal Colon (DC, red). **C)** Enrichment in differentiating epithelial cells (Colonocytes, Colonocyte-TA, Goblet, Goblet-TA). **D)** Enrichment in stromal cells (Telocytes, Interstitial Cells, Trophocytes). **E)** *Pdia3* and *Rpn1* (ER protein folding), as well as *Aifm2* and *Fancc* (p53 signaling) show rhythmic expression in telocytes. **F)** *Pip4p2* and *Itpk1* show rhythmic expression in Trophocytes. **G)** Enrichment in immune system cells.

We noted that many epithelial cells show enrichment for rhythms in endoplasmic reticulum protein folding genes (Fig 5A, 5C and Supplementary Fig 6A), and this rhythmicity is also present in the cells of the stroma including telocytes, trophocytes, and interstitial cells (Fig 5D). For example, endoplasmic reticulum associated genes such as *Pdia3* and *Rpn1* are rhythmic in telocytes especially in distal regenerating regions (Fig 5E). This highlights that protein processing is under 24-hour timing in different intestinal cells during regeneration, and it is notable that both epithelial and stromal cells are directly adjacent to the lumen where nutrients and microbiota are present. In addition to these common pathways, the stroma also exhibits cell-specific rhythms: telocytes show rhythms in p53 signaling (Fig 5D), that includes the genes *Aifm2* and *Fancc* (Fig 5E), while trophocytes show rhythms in phosphatidylinositol signaling that includes the genes *Itpk1* and *Pip4p2* (Fig 5F). Surrounding muscle cells MM and MP (Supplementary Fig 6B) are enriched in endoplasmic reticulum stress response, autophagy pathways, and FoxO / Insulin signaling. Indeed, rhythmically expressed signaling pathway genes are apparent in many different colon cells. The endothelium also exhibits enrichment for Insulin signaling as well as Integrin signaling (Supplementary Fig 6C); lymphatic cells show enrichment of Mapk signaling (Supplementary Fig 6D); ICC show enrichment of Tgfβ signaling (Supplementary Fig 6E), a pathway previously implicated in circadian phase timing^64^, with genes such as *Smad4* and *Tgfbr3* showing rhythmicity (Supplementary Fig 6F). These data reveal 24-hour mRNA rhythms in different cellular signaling pathways specific to different cell types.

Immune system cells were readily detected throughout the colon (Fig 3C-D), hence we examined rhythmic gene expression in these. Macrophages exhibit rhythms in many genes related to N-linked glycosylation and cholesterol metabolism (Fig 5G), whereas dendritic cells exhibit rhythms in cell cycle and Rho GTPase signaling (Fig 5G, Supplementary Table 1). This demonstrates that distinct processes have 24-hour rhythms in these innate immune system cell subtypes. Adaptive immune system cells that are very abundant in the colon show different mRNA rhythms. T-cell precursors (T-proliferative) and T-cells both exhibit rhythms in mRNA processing and regulation genes, while B-cell precursors exhibit rhythms in infection response and aerobic respiration (Fig 5G). Overall, the innate and adaptive immune system cells exhibit distinct 24-hour rhythmic transcriptional profiles. These data provide insight into the daily gene expression rhythms for colon-associated immune system cells. Our data provide a resource for future studies to assign specific gene rhythms to specific cells of the colon in undamaged and regenerating conditions (Supplementary Table 1).

### The epithelium exhibits lower amplitude circadian clock gene rhythmicity compared to surrounding tissue cells

Are these rhythmic genes in the colon related to canonical circadian clock activity? To test this, we examined the expression of clock components in a cell-type specific manner. Clock genes and clock target genes are widely expressed in various tissue compartments of the colon, with genes such as *Clock*, *Bmal1*, *Per1-3*, *Cry1-2*, *Dbp*, and *Tef* showing rhythmic 24-hour patterns (Fig 6A, Supplementary Table 1). This suggests the clock is functioning in this tissue, consistent with the free-running rhythms exhibited by colon explants (Fig 2J). However, the phase and amplitude of these genes differ between different cell types. While the muscle and stromal cells show similar timing and amplitude expression rhythms between proximal and regenerating distal colon regions, colonocytes, B-cells, T-cells, ISCs, and ICCs show lower amplitude in the same circadian clock transcripts in the proximal *vs*. distal regions (Fig 6A). In addition, the phases of clock components and clock target gene rhythms in epithelial cells are timed differently in some cases: for example, transcripts such as *Bmal1* and *Dbp* are anti-phasic in epithelia relative to the surrounding stromal compartment. This suggests the circadian clock transcription system is not co-phasic throughout the colon: different cell types exhibit different circadian clock gene timing.

**Figure 6.**
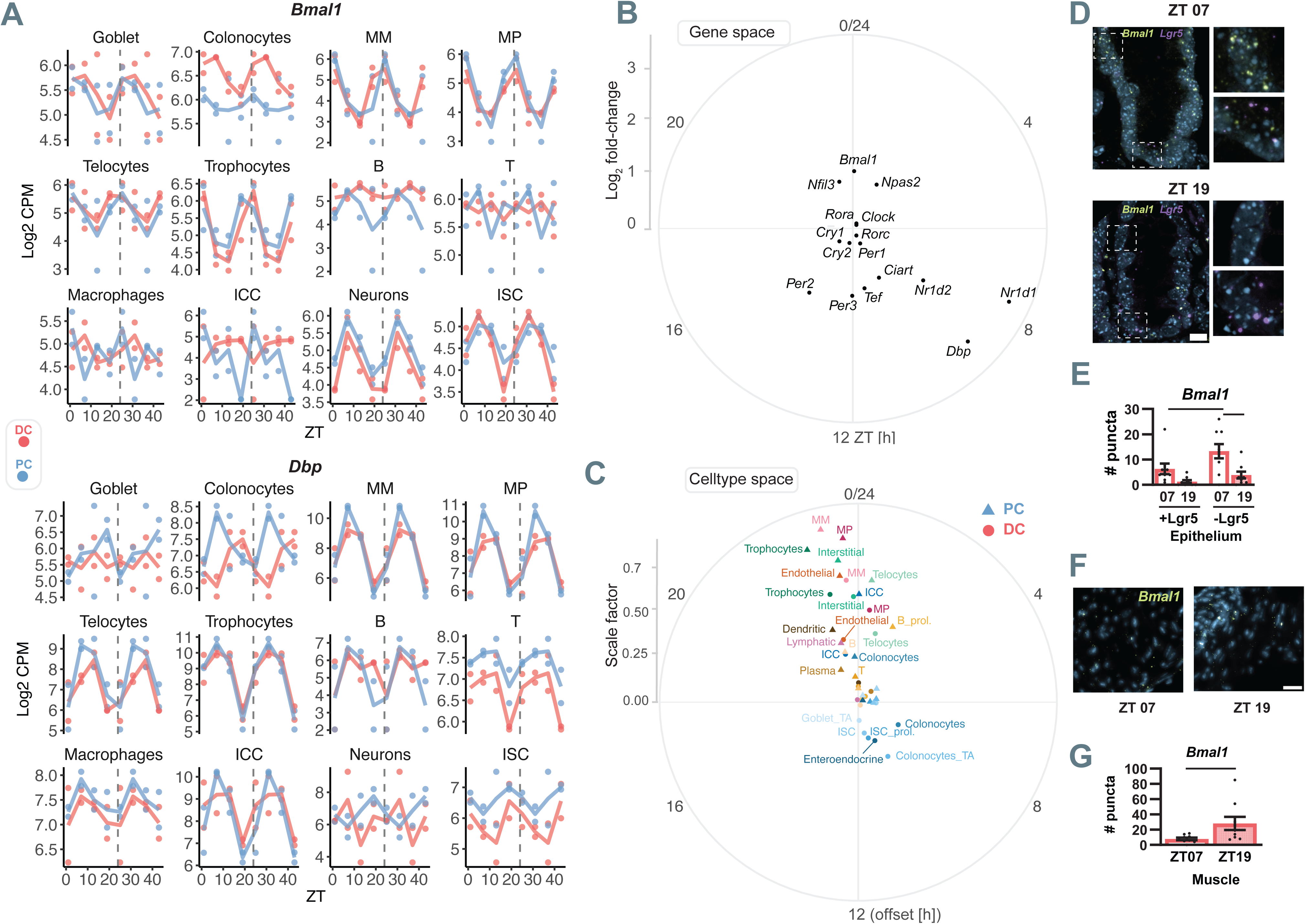
The epithelium exhibits lower amplitude circadian clock gene rhythmicity compared to surrounding tissue cells. **A)** Temporal normalized gene expression (Log_2_ CPM) of the core clock genes *Bmal1* and *Dbp* across several colon cell types. Dots represent the pseudo-bulk normalized count for each individual animal at a given ZT, and the colored line shows the average expression trend for the Proximal Colon (PC, blue) and Distal Colon (DC, red). Note that cells exhibit differences in the amplitude and timing of the same genes in a region and cell-specific manner. **B)** Gene Space: complex singular value decomposition (cSVD) analysis of clock genes across cell-types and regions. Polar plot shows the first gene complex (eigen-) vector (SVD1) from the cSVD performed on 12 canonical clock genes. The position of each gene represents its SVD1 phase (angle) and its Log_2_ fold-change (radius), reflecting the coordinated phase and amplitude of the core clock machinery. **C)** Celltype Space: polar plot shows the first cell-type complex (eigen-) vector (SVD1) from the cSVD analysis of canonical genes. The position of each cell type / condition represents the respective shift (angle) and scale factor (radius) of the gene module found in (B). PC samples are marked by circles and DC samples by triangles. **D)** Representative confocal images of *in situ* hybridization showing *Bmal1* and *Lgr5* expression at ZT07 and ZT19 in the distal colon during regeneration. Scale bar = 10µm. **E)** Puncta are increased at ZT07 in the distal colon epithelium. Puncta ware quantified in *Lgr5*+ and *Lgr5*-cells at each time point. Statistical analysis was performed using two-way ANOVA with Fisher’s LSD post hoc test. **F)** Representative *in situ* hybridization images showing *Bmal1* expression in the muscle. Scale Bar = 25 µm. **G)** Quantification of *Bmal1* puncta in muscle tissue at ZT07 and ZT19 in the distal colon. *Bmal1* expression is antiphasic between epithelium and muscle, with higher expression in muscle at ZT19. Mann–Whitney test was used. Significant differences (*p* < 0.05) are indicated by black bars.

To examine clock gene timing across cell types and regions, we summarized their 24-hour amplitude and phase relationships using complex-valued singular value decomposition (cSVD)^65^ (Fig. 6B). The dominant clock gene module extracted by cSVD reflects the coordinated timing of circadian genes across the colon. This module shows the typical rhythmic timing that would be expected of the clock system: *Bmal1* and *Npas2* peak in the morning, while *Nr1d1* and *Dbp* peak later in the day. *Tef* and *Per1-3* peak in the early evening, anti-phasic to *Bmal1,* followed by *Cry1-2*. The overall timing of these genes is as would be expected based on current knowledge of the clock, which is consistent with clock and clock target gene timing observed in previous bulk RNA-seq analyses of the intestine and colon^13,36,37,39–42^.

To examine how clock gene rhythms vary across cell types and regions, we visualized the right eigen-vector from the cSVD, which describes how the dominant gene module is expressed in each cell-type and region (Fig 6C). In these circular plots, the angle represents the phase relative (phase offset) to the dominant gene module, and the distance from the center indicates rhythm amplitude. Many colon cells exhibit robust and canonical clock timing including stromal cells (trophocytes, interstitial, telocytes), muscle (MM and MP), endothelium, and lymphatic cells. These cells also share clock phases relative to one another, withing a timeframe of ∼1-2 hours. Immune system cells are also within these times however with much lower overall amplitude, indicating lower clock and clock target gene rhythmicity. In most cases, distal cell types show lower amplitude rhythms compared to their proximal counterparts. For instance, trophocytes, interstitial cells, and telocytes have stronger rhythms in the proximal *vs*. distal regions, suggesting that regeneration dampens the clock gene expression rhythms. Unexpectedly, we discovered that the epithelial clock is distinct from the cells surrounding this tissue. Epithelial cells exhibit relatively weak circadian clock rhythms in the proximal colon, most of these show very low amplitude rhythms and thus cluster in the center. However, in the distal regenerating colon, epithelial cells show higher amplitude but cluster opposite with a timing that indicates the phase of rhythms is antiphasic or inverted by ∼12-hours. This is also clear when specific clock and clock target genes such as *Bmal1*, *Nr1d1*, and *Dbp* are examined: most colon tissue cells exhibit canonical clock gene expression timing, but epithelial cells do not (Supplementary Fig 7A-C). Indeed, not only is the epithelium antiphasic, but it also exhibits distorted phase coherence timing of clock and clock target genes both in the undamaged proximal and distal regenerating colon (Supplementary Fig 7D-F).

To further test epithelial clock timing, we probed for the gene *Bmal1* by fluorescence *in situ* hybridization (FISH) (Fig 6D). *Bmal1* was readily detected in the epithelial cells of colonic crypts at ZT07, the peak of its expression by scRNA-seq, and this was decreased at ZT19 when expression is lower (Fig 6A). To specifically probe for *Bmal1* expression in ISCs we co-stained for the gene *Lgr5*, a marker of ISCs^66^ (Fig 6D). Both Lgr5+ ISCs and Lgr5-epithelia express *Bmal1* higher at ZT07 than ZT19 (Fig 6D-E), suggesting this timing is present in both undifferentiated and differentiated epithelial cells, consistent with our scRNA-seq data (Fig 6A-C). In contrast, the surrounding muscle in the same tissue samples expresses *Bmal1* at higher levels at ZT19 than at ZT07 (Fig 6F-G), again consistent with the scRNA-seq analysis (Fig 6A-C). Together, these data indicate that the colon exhibits daily rhythms in circadian clock gene expression, but that epithelial cells and ISCs either show low clock gene oscillation (proximal colon) or exhibit a timing opposite to surrounding cell types in a regeneration-dependent manner (distal colon).

### Thermal response genes exhibit strong daily rhythms in the epithelium

The atypical expression of circadian clock gene rhythms in the regenerating epithelia seems inconsistent with the high number of 24-hour rhythmic transcripts detected in these cells (Fig 4A-B). What are the underlying factors influencing the expression of genes in the regenerating epithelium? To test this, we examined the top 200 pan-rhythmic genes, that show significant rhythmicity across the largest number of cell-type / region combinations and summarized their amplitude and phase relationships using the same cSVD approach as for the clock genes (Fig 7A-B). We noted that multiple heat shock proteins (Hsp) and associated chaperone protein coding genes such as *Hspa8*, *Hspa5*, *Hsp90b1*, *Hyou1*, *Fkbp4*, *etc*, peak in the early night at ZT12-16 when nocturnal mice are most active and have higher body temperature^67,68^(Fig 7B). At this time mitochondrial genes such as *Chchd2* and *Ndufa8* are also highest (Fig 7A-B), as well as immune system genes such as *Nlrp6*, *Pigr*, and *Dmpt1*. This may indicate a metabolic response to luminal nutrients and microbial factors that increase when food is consumed in a 24-hour rhythm. Twelve hours opposite to this timing, *cold inducible RNA binding protein* (*Cirbp*), a gene previously implicated in synchronizing the circadian clock to body temperature cycles^69–71^, reaches the peak of its expression (Fig 7A-B). Together the timing of the expression of these genes suggests that many rhythmic transcripts are responsive to physiological cycles driven by metabolic and microbial activity as well as body temperature during regeneration.

**Figure 7.**
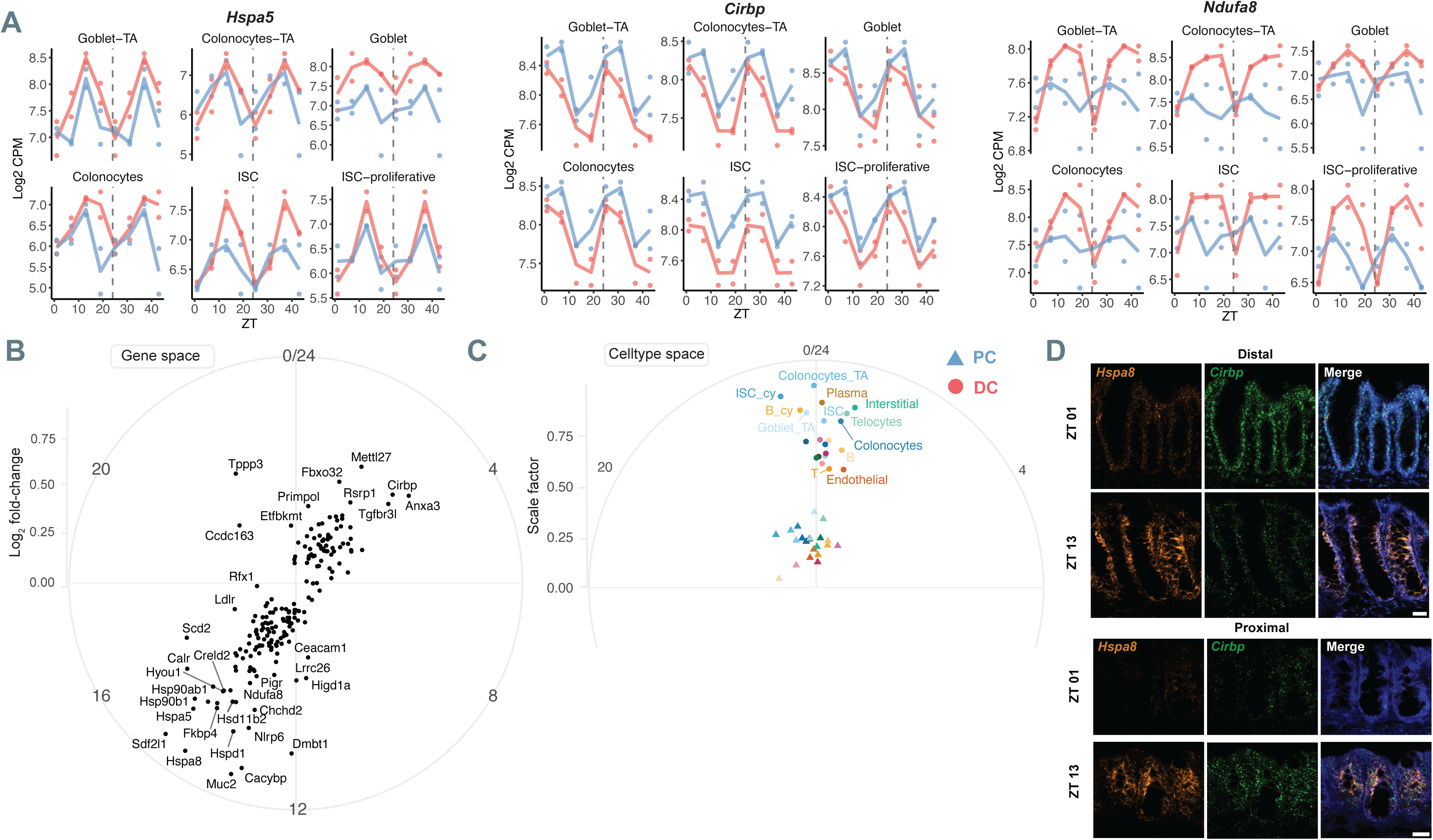
Thermal response genes exhibit strong daily rhythms in the epithelium. **A)** Temporal normalized gene expression (Log_2_ CPM) of *Hspa8*, *Cirbp*, and *Ndufa6* in selected epithelial cells. Dots represent the pseudo-bulk normalized count for each individual animal at a given ZT, and the colored line shows the average expression trend for the Proximal Colon (PC, blue) and Distal Colon (DC, red). **B)** Gene space: complex singular value decomposition (cSVD) analysis of clock genes across cell-types and regions. Polar plot shows the first gene complex (eigen-)vector (SVD1) from the cSVD performed on the top 200 pan-rhythmic genes across all cell types and regions. The position of each gene represents its SVD1 phase (angle) and its Log2 fold-change (radius), reflecting the coordinated phase and amplitude of the core clock machinery. **C)** Cell-type space: polar plot shows the first cell-type complex (eigen-)vector (SVD1) from the cSVD analysis of pan-rhythmic genes. The position of each cell type x condition groups represent the respective shift (angle) and scale factor (radius) of the gene module found in b) in the depicted sample conditions. PC samples are marked by circles and DC samples by triangles. **D)** Representative confocal images of *in situ* hybridization showing *Hspa8* and *Cirbp* expression at ZT01 and ZT13 in the proximal (scale bar= 20µm) and distal (scale bar= 25µm) colon during regeneration. *Hspa8* expression peaks at ZT13 in both regions, while *Cirbp* shows rhythmic expression only in the distal colon, peaking at ZT01 in antiphase to *Hspa8*.

We constructed cSVD plots showing cell-specific rhythms relative to the overall average (Fig 7C). As in Fig 6C, a time of 0/24h indicates the same gene timing in these transcripts in a cell type relative to the average and the amplitude shows the overall strength of this rhythmic structure. Distal colon epithelial, stromal, and immune system cells show a timing consistent with these cycles, suggesting specific cell types in regenerating regions exhibit a burst of gene expression synchronized with temperature and metabolic cycles. In contrast, most cells of the proximal undamaged colon do not have strong rhythms in these genes, indeed, these cluster in the centre, indicating low amplitude. The rhythmic expression of temperature, metabolic, and microbiome responses in the distal regenerating colon is consistent with its increased number of rhythmic transcripts (Fig 4A) and genes associated with feeding-driven pathways (Fig 4D).

To confirm the opposing timing of temperature response genes in the epithelium where this rhythmic structure is most apparent, we performed FISH for *Hspa8* and *Cirbp* to compare the peaks and troughs of these genes relative to one another (Fig 7D). *Hspa8* and *Cirbp* are highly expressed in epithelial cells relative to the surrounding tissue, and we note a slightly higher expression of these in the upper portion of distal crypts where differentiated epithelial cells are located (Supplementary Fig 7A-D). *Hspa8* is higher at ZT13 when mice are feeding and metabolically active, and lower at ZT01 when they are resting (Fig 7C-D, Supplementary Fig 7). In contrast, *Cirbp* is higher at ZT01 when body temperature is low, and lower at ZT13 when body temperature is high. The changes in these chaperone and heat-responsive transcripts suggests these cells are responding to proteome stress from circadian temperature and feeding cycles. This difference is less pronounced in the undamaged proximal colon, consistent with its lower rhythmic signature in these same genes (Fig 7B). These data show that the distal regenerating colon epithelium displays 24-hour rhythms in RNA transcripts responding to systemic factors.

## Discussion

Our data provide a chrono-atlas showing the cellular identity of daily rhythmic transcripts throughout the colon, including proximal regions that are undamaged and distal regions that are undergoing regeneration post-injury. Most of the cells in the intestine (*i.e.* stroma, muscle, endothelium, immune system) have circadian clock gene rhythms timed as we expect from a functional clock, indicating the circadian timekeeper functions in these cells. Undamaged proximal colon cells appear to show stronger circadian clock rhythms that are largely in phase in most cell types within a ∼1-hour timing (Fig 6C). Distal regenerating colon cells show lower amplitude clock gene rhythms relative to their proximal counterparts (Fig 6C), however circadian clock rhythmicity mostly persists in these cells during regeneration (Fig 2). In addition to clock genes, we find thousands of genes showing 24-hour rhythms in both proximal and distal colon regions (Fig 4A). As the mice sampled in these experiments were housed in light-dark conditions, we cannot resolve whether these genes are driven by the endogenous circadian clock present in colon cells or reflect responses to rhythmic systemic signals such as daily rhythms in physiology (e.g. body temperature), feeding, or microbial activity. Indeed, our findings that the regenerating colon exhibits transcript rhythms in a biphasic pattern, with peaks at the transitions between active / inactive times of day (Fig 4B) and shows a strong thermal response (Fig 7), suggests many gene rhythms are elicited outside of the colon itself. These daily systemic cues influencing colon rhythms are downstream of behaviour and physiology that is itself driven by a circadian clock located in the brain^72^. The time of feeding has emerged as a potent synchronizing cue in digestive tissues such as the liver and gastrointestinal tract^23,42,73,74^, hence it is expected that the colon would display transcriptional waves driven by nutrients and microbial metabolites that arise when food is consumed. Our data provide a framework for future studies of colon physiology by shedding light on the 24-hour timing of specific cell types in this complex and dynamically functioning tissue. Because DSS injury is a widely used laboratory model of colitis, these data are relevant to understanding the daily timing occurring during pathological states relevant to IBD and its resolution through regeneration.

Relative to the other cells of the colon, the epithelium has either weak (proximal undamaged) or anti-phasic (distal regenerating) clock and clock target gene rhythms (Fig 6, Supplementary Fig 7). Recent studies using microarrays or bulk RNA-seq have documented clock gene rhythms in the gastrointestinal tract^13,36,37,39–42^, and conditional loss of *Bmal1* in the epithelium has been shown to affect colitis outcomes^30–34^. This has been understood to mean that the epithelial circadian clock is functional in generating daily transcriptional rhythms and that the loss of these, through the required clock gene *Bmal1*, is detrimental^75^. Our results thus reveal unexpected features of epithelial clock timing relevant to these studies. First, the general clock structure detected in previous bulk genomics studies most likely reflects transcripts detected from the surrounding support cells (*i.e*. stroma, muscle) and to a lesser extent from the epithelium (Fig 6). The weak or anti-phasic rhythms in clock gene rhythms from the epithelium are obscured at the whole tissue level and were previously unappreciated. Indeed, it is not clear whether the circadian rhythmicity measured in *Per2-Luc* colon explants *ex vivo* reflects the anti-phasic clock of the epithelium or the signals emanating from the underlying support cells underneath (Fig 2J). However, our results are not incompatible with an epithelial clock transcriptional timer but rather reveal that this timer is not in phase with the overall clock rhythms that have been previously detected by bulk RNA-seq. The anti-phasic epithelium clock in the regenerating distal region is likely to drive a transcriptional program, and we note that many daily rhythms are present in the epithelial cells that include both clock and response-driven targets (Fig 4D-E, Fig 5, Supplementary Table 1). Second, the observation that the conditional loss of *Bmal1* generates a phenotype following DSS administration could be the result of the loss of the epithelial clock transcription rhythm or non-circadian roles of *Bmal1* in the epithelium. We note that discrepancies have been found in *Bmal1* epithelial loss of function experiments, generating an increased^37^ *vs.* decreased^76^ pathology, *vs.* no pathology^77^ following DSS-induced colitis. The low amplitude clock structure of the normal epithelium, coupled with its enrichment for metabolic and stress response gene rhythms (Fig 5, Fig 7), suggest that environmental factors play a strong role in shaping epithelial daily rhythmicity. It may be the case that the cell-autonomous epithelial clock is subordinate or masked by daily extrinsic gene rhythms arising from behaviour that drives daily changes in physiological states. It is thus not clear how the loss of epithelial clock function relates to overall tissue transcriptional rhythms, and their subsequent function and pathology.

It is surprising nonetheless that the regenerating epithelium can display atypical clock gene rhythms relative to the surrounding stromal telocytes, trophocytes, and interstitial cells that reside in direct contact with the epithelial cells (Fig 6). How would this be achieved? One possibility is that the cues synchronizing the clocks across cells in the epithelium *vs.* stroma are different. The epithelial clock could be subject to cues such as body temperature, nutrients, and microbial metabolites while the stromal clock may be responsive to hormones, and cytokines. The integration of different Zeitgebers (time entrainment factors) in these different cell types could lead to different circadian clock timing. An enrichment of metabolic and nutrient response pathways in the epithelium but not the stroma supports this possibility (Fig 4D, Fig 5A-F). We also note that along with the epithelium, many cells of the immune system also have weak clock rhythms in the colon (Fig 6C), suggesting that certain cells that are directly exposed to luminal nutrient and microbial factors may have disruptions in circadian clock rhythms. Food intake may resynchronize the clocks of cells directly exposed to food- or microbiome-derived chemicals. It is indeed possible that in the context of regeneration, weak clock gene rhythms in the epithelium now become highly sensitive to proteome stress driven by daily environmental factors, that these become offset in time by 12-hours. Future studies will resolve the mechanism underlying this anti-phasic timing between adjacent cells in the regenerating intestine. At present, we note that the finding that adjacent cells in an organ have offset timing is not without precedent. In the central suprachiasmatic nucleus of the brain, neuronal clocks are a few hours earlier than in the adjacent glia^78^. This circadian clock phase disparity in adjacent cells is thought to arise from inhibitory neuron-to-astrocyte signaling that slows the astrocyte clock. It is possible that a similar inhibitory signaling relationship may be occurring in the epithelium and stromal cell lineages in the intestine.

Both regions of the colon that we tested display rhythms in thousands of transcripts (Fig 4A). Rhythmic genes in the proximal colon can peak throughout the day with an increase at midday (Fig 4C) and an enrichment in many cell signaling, proliferation, and growth-related pathways (Fig 4E). In contrast, we find that the regenerating distal colon predominantly shows biphasic rhythms with morning and evening peaks of gene expression (Fig 4B) corresponding to pathways involved in metabolism, nutrient responses, stress responses, and protein production / recycling (Fig 4D). Together, these observations are consistent with other studies of the colon using bulk RNA-seq that have found enrichment in metabolism, cell signaling, and microbial response rhythms^13,37,39^. Similarly in the small intestine, bulk RNA-seq has shown that there are rhythms in transcripts encoding cell growth, protein production, and metabolic responses^14,36,40,41^. While these bulk data provided averaged mRNA signals, our data now provide further resolution to these studies, identifying regional differences that are related to regeneration state and the precise cellular source of these rhythmic genes. Of particular interest in the field has been the effect of the microbiome in shaping intestinal gene expression and how this is related to daily cycles of food consumption. Many of the rhythms found in our study are likely to depend on daily rhythms in microbiome-derived metabolites, as metabolic and stress responses have been noted to be absent in germ-free or antibiotic conditions^13,36,37,41^. Future studies carried out on the colon under germ-free conditions would resolve which rhythmic genes are driven by microbial factors.

It has been long noted that the intestine and colon exhibit rhythms in epithelial cell proliferation^79,80^, rhythms in the transcription of cell cycle regulators^38,81^ or epithelial signaling pathway genes^39^, and that loss of clock genes abolishes these processes^30,32,82^. As in the context of colitis, this has been understood to mean that circadian clock regulation of gene expression rhythms drives the ability of ISCs and precursor cells to divide or differentiate according to a 24-hour cycle. Here, we observe rhythms in genes related to Notch signaling in ISCs (Fig 5A), Egf signaling in Tuft cells (Supplementary Fig 6A), and Hippo signaling in colonocytes (Fig 4E). These are consistent with this model, and we do note that cell cycle regulators are present in both proximal and distal colon cells (Fig 4D-E, Fig 5A). However, signaling genes do not seem to be enriched in the distal regenerating regions where the intestinal epithelium is undergoing the most active proliferation. Instead, in the regenerating colon we note high enrichment in genes involved in metabolism (oxidative phosphorylation, citrate cycle, purine metabolism, *etc*.) and protein production (protein processing in endoplasmic reticulum, autophagy, *etc*.) (Fig 4D). These appear to be shared by multiple cell types of the colon including ISCs and other epithelia, stroma, immune system, and muscle. Hsp are highly enriched among these genes, notably, a recent paper has found that Hsp proteins regulate ISC survival and epithelial barrier function thus providing a protective effect during DSS-induced colitis^83^. Could rhythms in these processes explain rhythms in colon proliferation? It has also been shown that ISCs in the small intestine are enriched for a fatty acid oxidation program that drives their maintenance and proliferation^84,85^. Our results are consistent with these programs being elevated during regeneration and show that genes related to ISC metabolic function may be occurring with a 24-hour timing to drive proliferation cycles in colon cells. Recently it was shown that ISCs are highly sensitive to feeding following a period of fasting, which causes a rapid induction of mTORC1 activity and upregulation of protein synthesis^86^. This drives an ISC proliferative state when nutrients become available following a fast, another related nutrient-dependent mechanism that could couple with daily transcriptional cycles to account for a proliferation rhythm in the regenerating colon. A third possibility is the concept called paligenosis^87^, whereby differentiated cells undergo cellular remodelling through mTORC1 inhibition and upregulation of autophagy to dedifferentiate into stem cells. It is possible that the daily rhythms in the transcripts related to protein production, endoplasmic reticulum stress, and protein recycling that we observe are linked to paligenosis in dedifferentiating colon epithelia. Metabolic and protein-based daily rhythms provide a rich area of investigation for future studies to use this dataset as a resource to interrogate the connections between environmental factors and cellular timing.

## Limitations of the study

We sampled proximal and distal regions of the colon and inferred that regeneration is responsible for the changes in rhythmic programs that we observe between undamaged *vs.* regenerating regions. However, some of these differences may be due to anterior *vs.* posterior transcriptional differences; future studies sampling undamaged and regenerating tissues separately will resolve this issue. Due to cost and logistics, our sampling was limited to four time points over 24-hours and a cohort of 8 mice with both males and females represented equally. As such we may have missed detecting all rhythmic genes. Future studies with more time points tested, deeper sequencing with more cells analyzed, and larger cohorts of both male and female mice would improve this analysis. Together these additional tests would provide a more detailed list of all 24-hour rhythmic transcripts in a sex-specific manner.

## Methods

### Animal housing and breeding

*Per2-Luciferase* mice (Jackson Laboratories, *B6.129S6-Per2tm1Jt/J* strain #006852)^49^ were housed and bred at the University of Windsor animal facility. Littermates were co-housed under a light dark cycle (LD 12h:12h, light onset = ZT0, 250lux) with ad libitum access to food (PicoLab-Diet, Cat. No. 5053) and water. All experiments follow regulations and approved protocols by University of Windsor(AUPP 23-17).

### DSS administration

Mice (13-15 weeks old) were individually housed 14 days before the administration of 3.5% DSS (MW: 40 KDa, Boc Sciences, Cat no. 9011-18-1) in the drinking water for 7 days, followed by 10 days with regular water. Colitis symptoms were evaluated using Disease Activity Index (DAI) during DSS treatment^88^. Cage changes were performed near light offset (ZT12) on days D0 and D10 (Supplementary Fig.1a), transferring 20% of original bedding and nest to the new cage.

### Tissue collection

Colon tissue was collected at ZT01; before DSS (d0) and after DSS treatment (d7, d10, d17). Mice were euthanized using CO2 followed by cervical dislocation, colons were flushed with ice cold PBS (Corning, Cat no. 21-040-CM), open longitudinally and rolled from proximal to distal end for histological analysis. Tissues were fixed in 4%PFA in PBS for 2h room temperature, cryopreserved in 30% sucrose in PBS for 48h at 4°C and placed in cryo-embedding compound for storage at -80°C.

### Histology and immunohistochemistry

10µm sections were used for histological analysis. The total length of amorphous (crypt absent) lesions, and hyperplastic crypt regions (regenerative) were quantified from Hematoxylin & Eosin-stained sections. Cell proliferation was assessed from Ki67 immunolabeled sections (1:100, Abcam, Cat no. ab64261) detected by HRP/DAB kit as previously described (Abcam, Cat no. ab64261)^30^. Percentage of positive cells were quantified in 6 crypts in proximal and distal region.

### Per2-Luciferase recording

For the ZT01 group, colon segments (400 µm × 400 µm pieces) from the most proximal and distal regions were cultured for bioluminescence recording over 4 days, as previously described (PMID: 35223537; PMID: 29054877).

### In situ hybridization and microscopy

Fluorescent in situ hybridization was performed using the RNAscope multiplex fluorescent detection kit (Advanced Cell Diagnostics, Cat no. 323280) with the following probes Mm-Hspa8 (Cat no. 113791-C2), Mm-Cirbp (Cat no.534421) at ZT01 and ZT13; and Mm-Lgr5 (Cat no.312171) and Mm-Arntl1 (Cat no.438241-C2) at ZT07 and ZT19. Images were acquired from 3 proximal and 6 distal regions. Signal was quantified from 5-6 cells per region. For *Hspa8* and *Cirbp*, total puncta and mean optic density were measured at the crypt top and base. *Bmal1* puncta were quantified in *Lgr5* positive and negative cells. In muscle tissue, puncta were counted within regions of 1700 µm^2^ area. Histological images were captured using Zeiss Axio Scan.Z1 slide scanner or Zeiss LSM900 confocal microscope. Image analysis was performed with Zen software (Zeiss) or QuPath software^89^.

### Single-cell RNA sample collection

Mice were individually housed in behavioral chambers (Actimetrics, Inc) two weeks before DSS treatment. Behavioral recordings began one week after individual housing and continued throughout the experiment (Supplementary Fig. 2a). Food intake events were recorded using an automatic feeder system that dispenses dustless precision pellets (315 ± 4 mg each containing 3.35 kcal/g, BioServ, Cat no. F0170). Food intake and free-running behavior were collected and analyzed with ClockLab software (Actimetrics Inc.)^90^. To minimize behavioral disruptions, disease activity index was assessed on days d5, d7, d10 and d17. On day 17 (Supplementary Fig. 2a), colon tissue was collected at 6-hour intervals (ZT01, ZT07, ZT13, ZT19, n=2 by ZT). The colon was divided in half; one half was rolled for histology; the other was divided into proximal and distal segments. After removing PBS, tissues were flash-frozen in liquid nitrogen. Single-cell RNA-seq libraries were prepared with 10X Genomics Flex Gene Expression assay (Princess Margaret Genomic Center, Toronto). Tissue samples were processed using the chop/fix method (Protocol CG000533 – Rev A) into fixed single cell/nuclei suspensions. Tissues were incubated for 25 minutes on the octo dissociator. Protocol CG000527 - Rev D was used to perform the Flex assay (probe barcode hybridization, GEM generation, GEM recovery & Pre-amplification, Library construction). Samples were sequenced on the NovaSeq 6000.

### scRNA-seq analysis

Raw reads were aligned to the mouse reference genome (GRCm39, refdata-gex-mm10-2020-A) and quantified using Cell Ranger multi (10x Genomics, cellranger version 7.0.1) with default parameters. The resulting count matrices were analyzed in *Seurat* (v4.1.3) in R. Low-quality cells and potential multiplets were removed based on the following thresholds: a minimum of 500 and maximum of 15,000 unique molecular identifiers (UMIs) per cell, detection of 500-7,500 genes per cell, and a maximum mitochondrial read fraction of 7.5%. After quality control, gene expression values were normalized and log-transformed, and highly variable genes were identified using the FindVariableFeatures function (*n* = 2,000; method = “vst”). To correct for batch effects and identify cell types, datasets were integrated using reciprocal PCA (rPCA) with default parameters. Principal component analysis (PCA) was performed on the integrated data, and the first 30 PCs were used to construct a shared nearest-neighbor graph. Clustering was performed using the Leiden algorithm at a resolution of 0.5. For visualization, cells were embedded in two dimensions using uniform manifold approximation and projection (UMAP) based on the same 30 PCs. Cell types were annotated using canonical marker genes from the literature. Fibroblasts, smooth muscle cells, and transit-amplifying (TA) colonocytes were further resolved by subclustering (FindSubCluster, resolution = 0.05) into subtypes based on marker gene expression (Supplementary Fig. 4).

Genes with fewer than 1,000 total counts across all cells were excluded. For downstream differential and rhythmicity analyses, pseudo-bulk profiles were generated by summing raw counts across all cells within each cell type, region, and animal. Pseudo-bulk count matrices were analyzed in *DESeq2* : counts were normalized using the trimmed mean of M-values (TMM) method and log_2_-transformed with a prior count of 2.

Rhythmicity was assessed using linear harmonic regression, on the log_2_-normalized count, of the form: 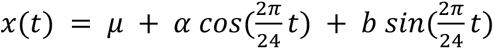. Amplitude was computed as 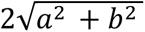 and phase as 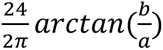. Statistical significance of rhythmicity was assessed by an R^2^-based test using the beta distribution. Prior to fitting harmonic regression, genes with insufficient expression were excluded to reduce false positives. Within each cell type and condition, genes with fewer than 50 total counts across all samples or expressed at fewer than five time points were not tested. In addition, cell types represented by fewer than 1,000 cells were excluded from rhythmicity analysis.

cSVD. Genes passing rhythmicity thresholds (p < 0.05, log2 amplitude > 0.5) were retained for Complex-valued singular value decomposition (cSVD); entries with p > 0.2 were set to zero to reduce uncertain estimates. For each cell type × condition, we constructed a complex matrix, decomposed it by cSVD, and post-hoc rotated/scaled components so that group loadings shared a common phase reference and gene loadings reflected singular values as in^65^. Amplitudes were visualized in polar coordinates (phase as angle, modulus as radius), and interpreted as joint phase-amplitude modes of coordinated rhythmic programs across groups. cSVD was applied either to canonical clock genes or to the top 200 pan-rhythmic genes across all cell types and conditions; only the first module (SVD1) is shown.

To identify the biological pathways under circadian control, gene set enrichment analysis was performed on the lists of genes determined to be rhythmically expressed in each colon cell type x regions. Rhythmic genes were submitted to enrichR using the KEGG_2019_Mouse, WikiPathways_2024_Mouse, and Reactome_Pathways_2024 databases. Pathways were retained for analysis only if they contained more than three overlapping rhythmic genes and fewer than 500 total genes. Terms related to disease or infection (e.g., “cancer,” “viral”) were excluded. For visualization, the top five most significantly enriched terms (Adjusted P-value < 0.1) for each cell type and database were selected. The final list was ordered by the minimum adjusted P-value across all groups.

### Statistical analysis

All the non-scRNA-seq reported experiments used three biological replicates, unless otherwise specified in the figure legend or main text. No samples were excluded from the analysis. All histological analyses were performed blinded. GraphPad Prism 9 was used for performing statistical analyses. All data are presented as mean ± s.e.m. Normality was assessed using the Shapiro–Wilk test, and parametric or nonparametric tests were selected accordingly. Specific statistical details are provided in the figure legends and in Supplementary Table 2.

---

**Supplementary Figure 1.**
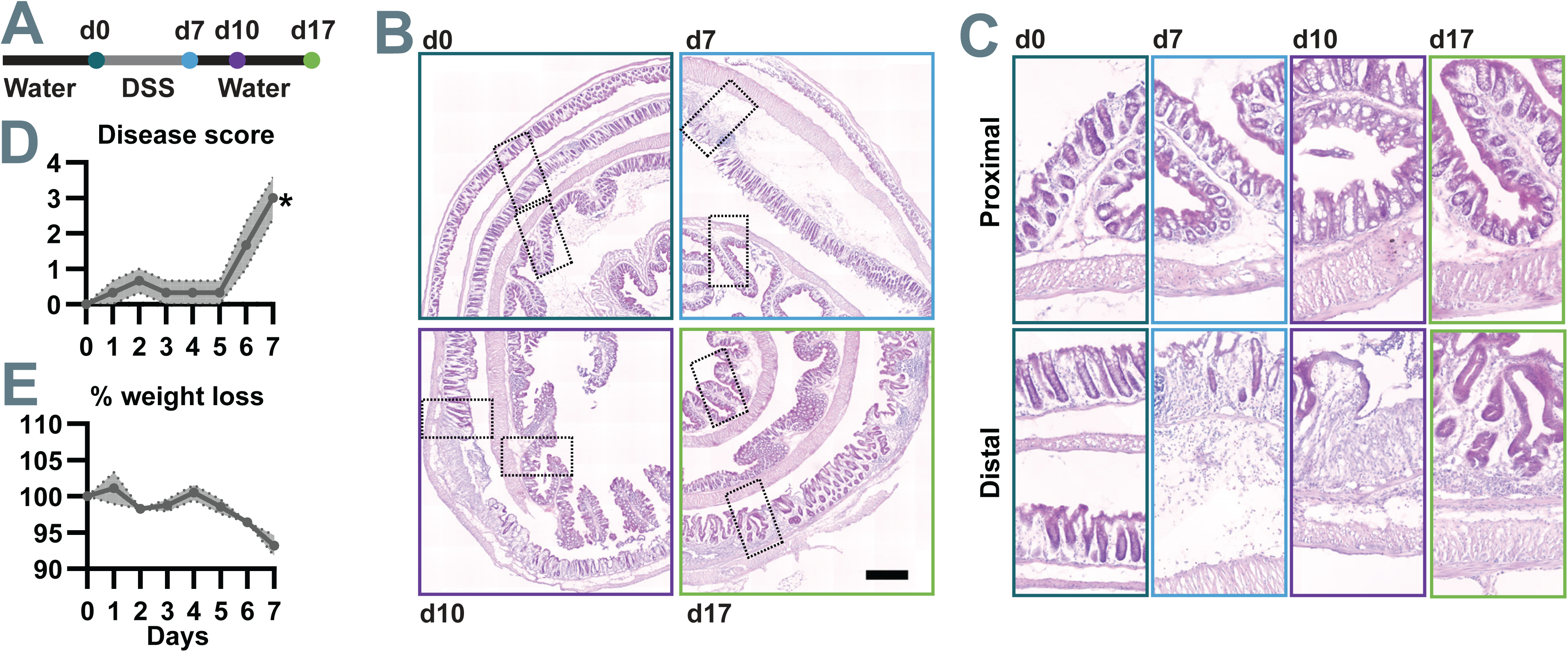
Disease progression and histopathology. **A)** Schematic of experimental timeline showing 7 days of 3.5% DSS administration and tissue collection. **B)** Disease score tested during DSS treatment indicates symptoms of damage are evident (Friedman test with Dunn’s post hoc test relative to day 0. **C)** Percentage of body weight lost during colitis (repeated measurements one-way ANOVA with Dunnett’s post hoc test relative to day 0. **D-E)** Representative Hematoxylin and Eosin-stained colon images at the indicated time points; scale bar = 500 µm. Dashed boxes indicate regions that are shown magnified in panel C. Data are presented as mean ± s.e.m. n=3 animals by time of collections. Asterisks indicate significant differences (p<0.05) relative to day 0.

**Supplementary Figure 2.**
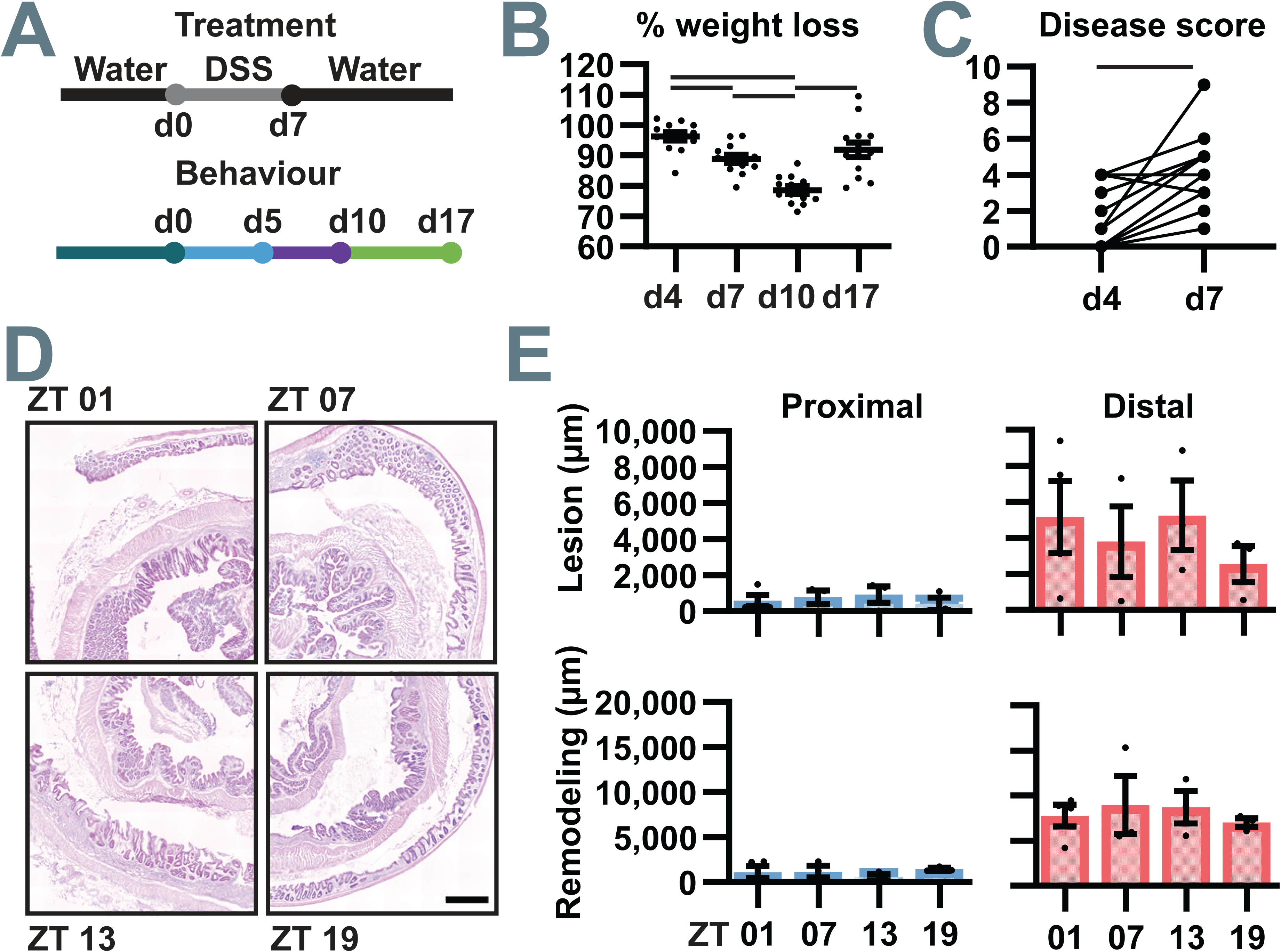
Analysis of samples shown in Fig 2: Circadian rhythms persist during regeneration. **A)** Schematic of experimental timeline showing days of DSS treatment as these correspond to periods of time analyzed for behavior. Color code indicates behavioral analysis carried out: dark blue = no damage, light blue = no disease symptoms, purple = symptoms of injury apparent; green = recovery phase. **B)** Percentage of body weight lost during DSS treatment peaks at day 10, with recovery in weight observed by day 17 (repeated-measurements one-way ANOVA with Tukey’s post hoc test) **C)** Disease symptom score increases by day 7, indicating worsening symptoms at the individual level (Wilcoxon test). **D)** Representative Hematoxylin and Eosin-stained sections from colons collected at different times of day (ZT01, ZT07, ZT13, ZT19) at day 17; scale bar = 500 µm. **E)** Quantification of total lesion length (upper graphs) and regenerating region size (lower graphs) across 24-hour shows no significant differences (Kruskal-Wallis test). Data are presented as mean ± s.e.m. n = 12 mice (n=3 per timepoint).

**Supplementary Figure 3.**
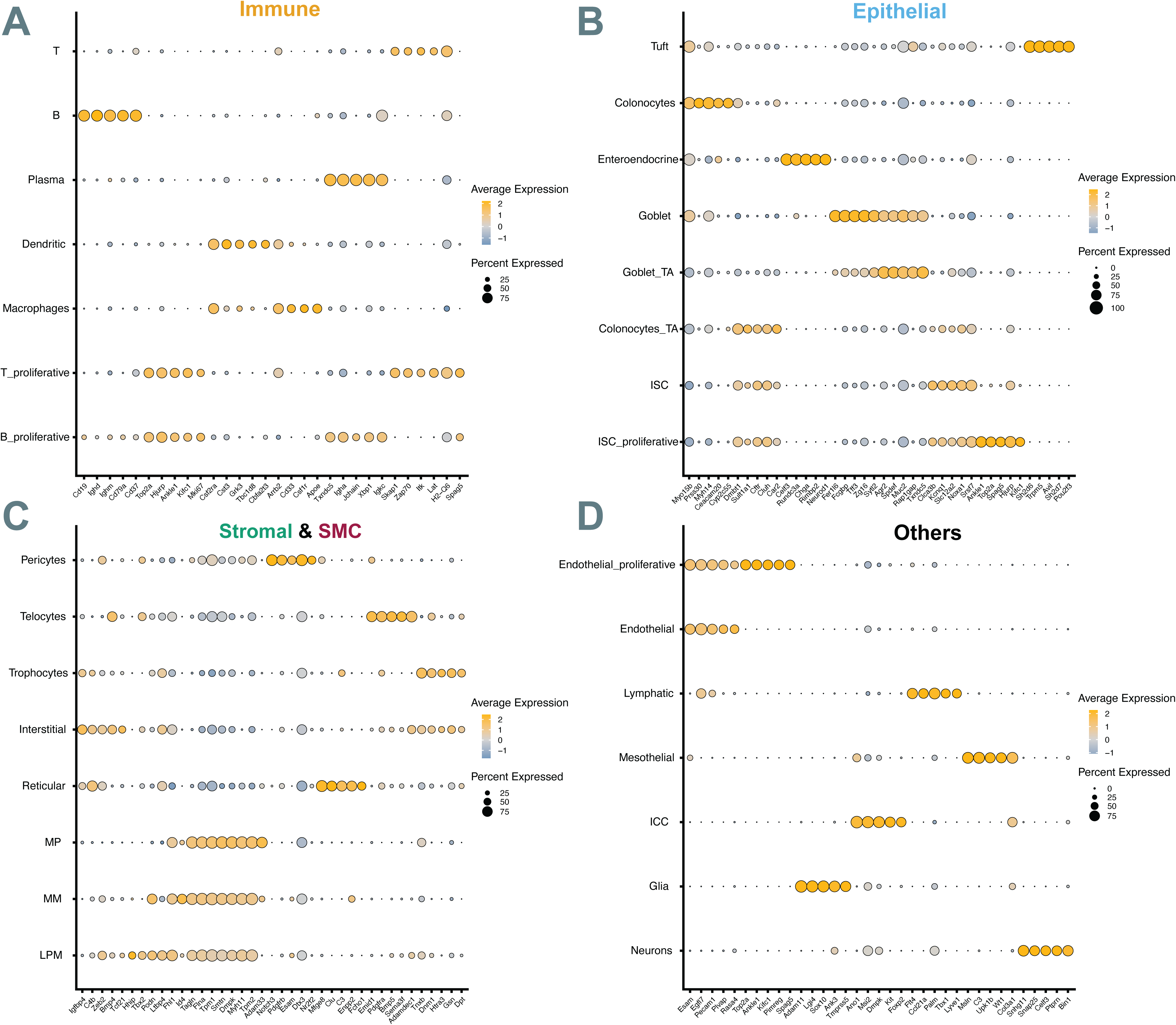
Cell type-specific marker gene expression across major cell populations. Dot plots showing discriminative marker genes for: **A)** Immune system cells. **B)** Epithelial cells. **C)** Stromal and smooth muscle cells (SMC). **D)** Other cell types. Dot size represents the percentage of cells expressing each gene within a cell type. Color intensity indicates average scaled expression level (z-score) across cell types, with yellow representing higher relative expression and blue/gray representing lower relative expression. Marker genes were identified using Wilcoxon rank-sum test within each broad category (n=5 top markers per cell type, AUC > 0.4, adjusted p-value < 0.05).

**Supplementary Figure 4.**
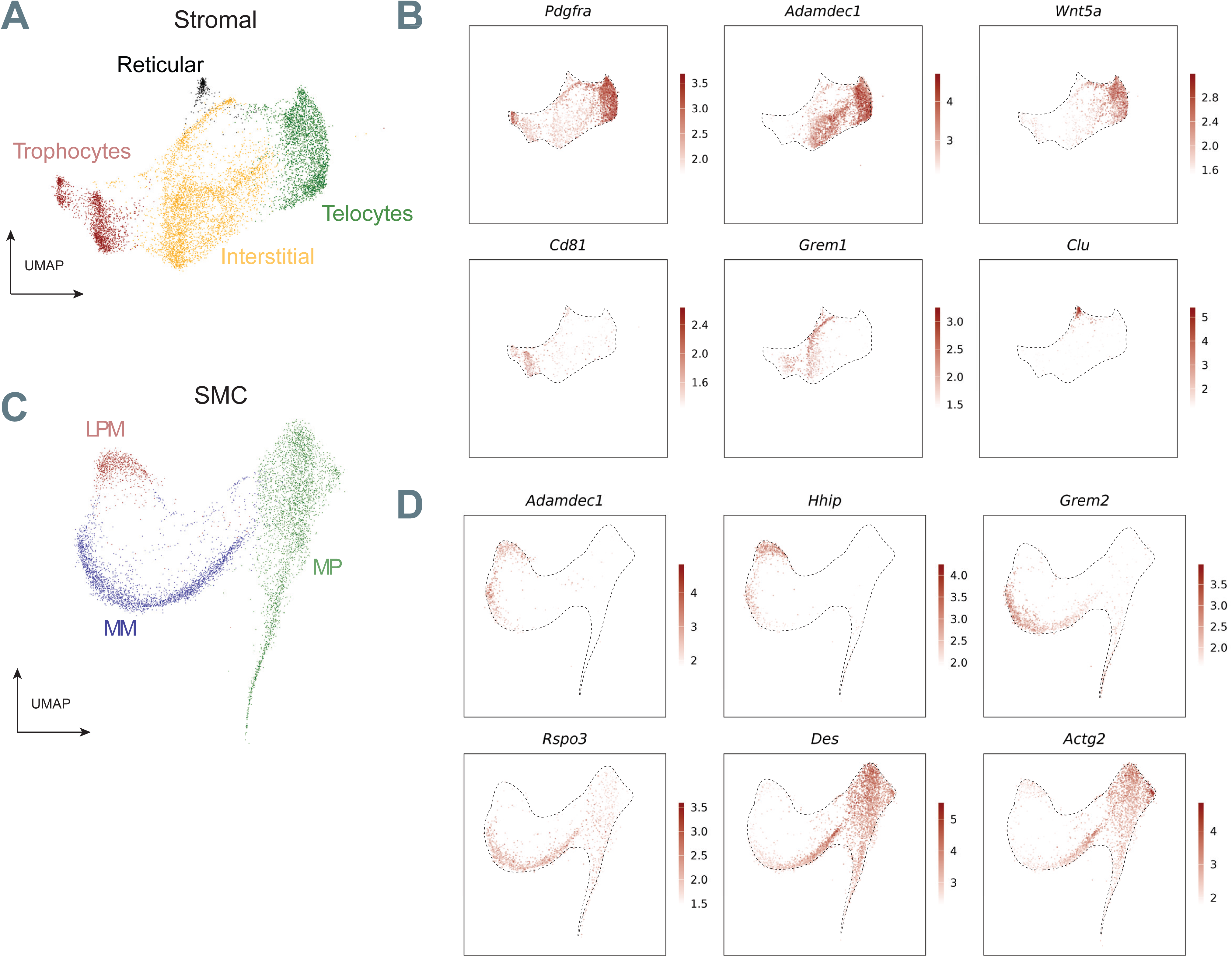
Sub-clustering of stromal and smooth muscle cell populations. **A)** UMAP visualization of stromal cell subclusters including trophocytes, interstitial cells, telocytes, and reticular cells. **B)** Feature plots showing expression of stromal marker genes in these subclusters (*Pdgfra, Adamdec1, Wnt5a, Cd81, Grem1, Clu*). **C)** UMAP visualization of smooth muscle cell (SMC) subclusters including lamina-propria-associated myocytes (LPM), muscularis mucosae (MM), and muscularis propria (MP). **D)** Feature plots showing expression of SMC and myofibroblast marker genes in these subclusters (*Adamdec1, Hhip, Grem2, Rspo3, Des, Actg2*). Color intensity represents normalized gene expression levels.

**Supplementary Figure 5.**
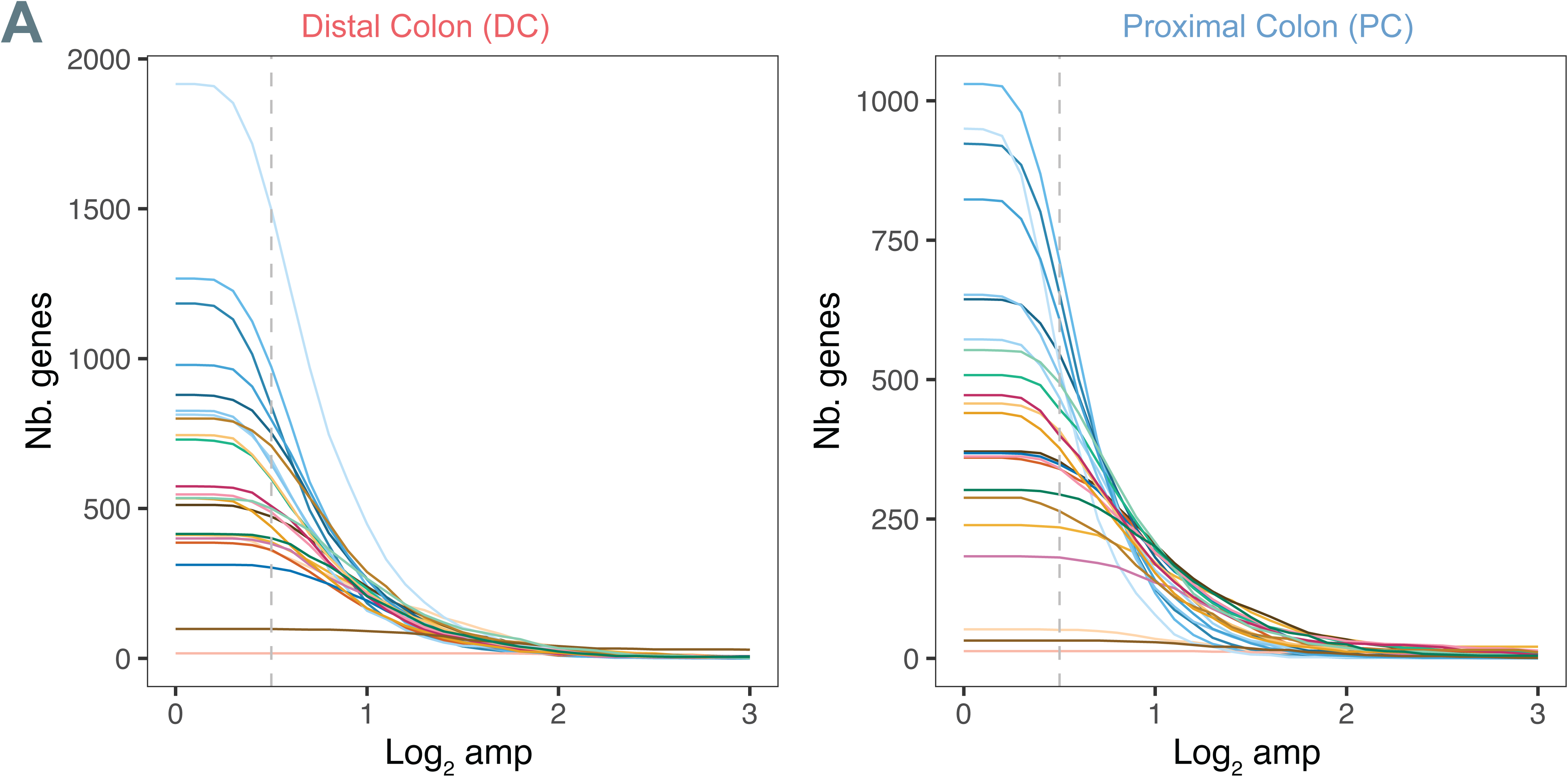
Number of rhythmic genes as a function of oscillation amplitude in the distal and proximal colon. **A)** Number of genes showing significant rhythmicity (p < 0.05, harmonic regression) in the different cell types, plotted as a function of increasing log_2_ peak-to-through amplitude. The dashed line (0.5 log₂ amplitude) indicates the amplitude threshold used in the following analyses to define a gene as rhythmic.

**Supplementary Figure 6.**
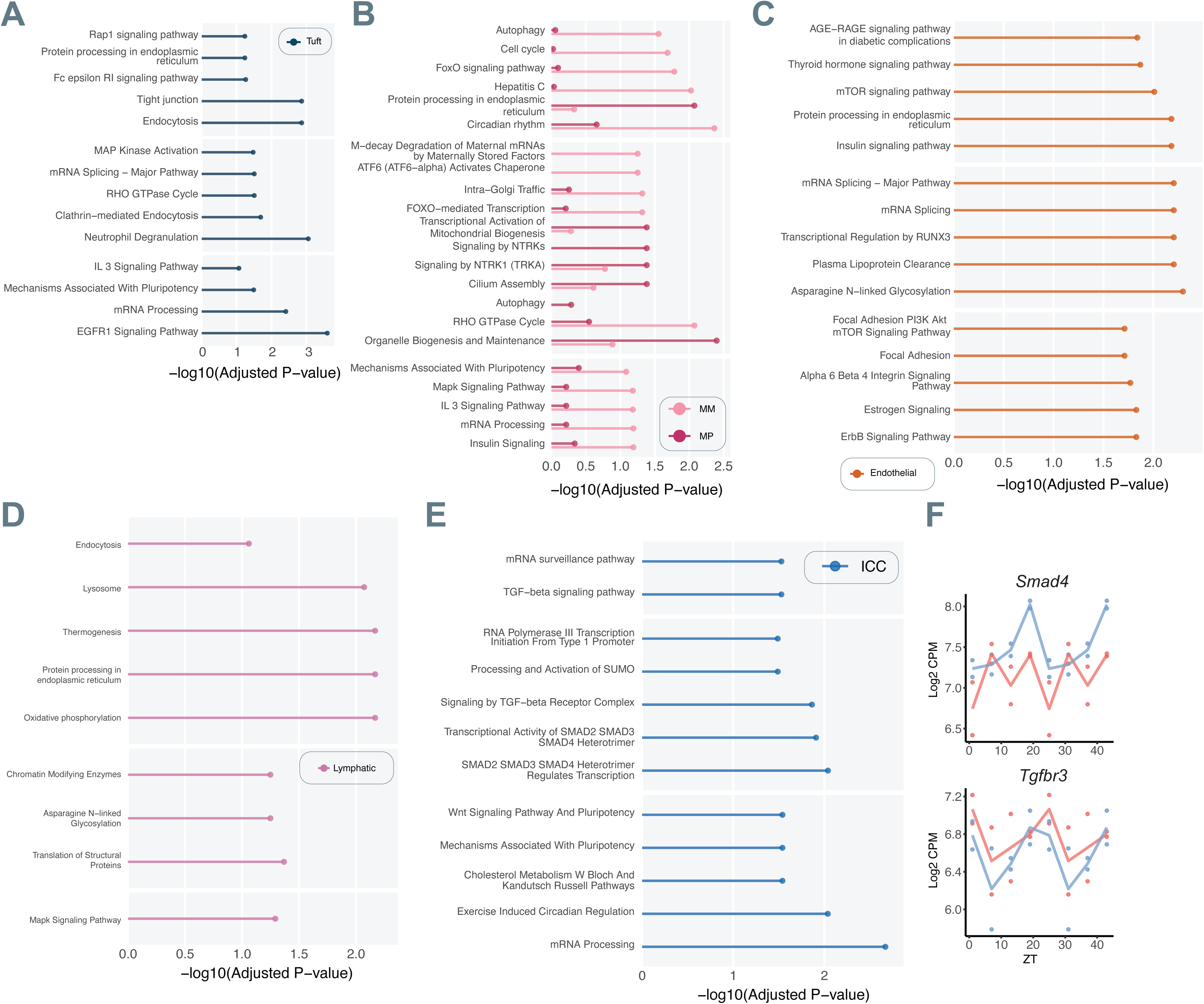
Cell-type specific 24-hour rhythms in gene expression across colon cell lineages. Enrichment of rhythmic genes in specific cell types for major biological pathways. Bar plots show the negative log_10_ of the adjusted P-value (-log_10_ adjusted p-value) from Enrichr analysis (KEGG, REACTOME, WikiPathways) for genes exhibiting 24-hour rhythms (p-value<0.05 and Log 2 peak-to-trough amplitude >0.5) in each specified cell type. **A)** Tuft cells. **B)** Muscularis Mucosae (MM) and Muscularis Propria (MP). **C)** Endothelial cells. **D)** Lymphatic cells. **E)** Interstitial cells of Cajal (ICC). **F)** *Smad4* and *Tgfbr3* normalized expression level (Log_2_ CPM). *Smad4* shows 24-hour rhythms the proximal colon (PC, blue) but not the regenerative distal colon (DC, red), while *Tgfbr3* shows rhythms in both. Dots represent the pseudo-bulk normalized count for each individual animal at a given ZT point, and the colored line shows the average expression trend.

**Supplementary Figure 7.**
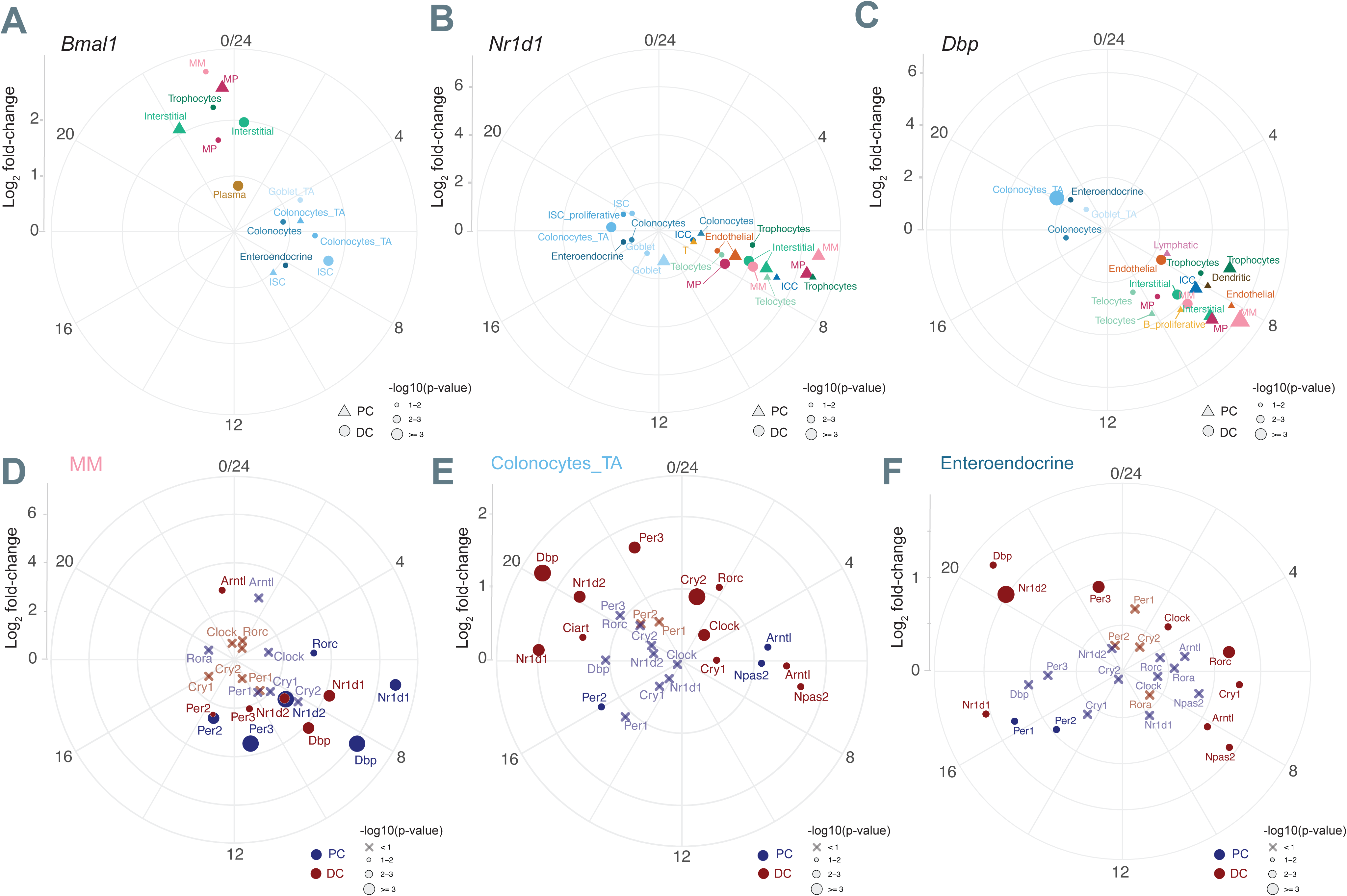
Core clock and clock output genes show antiphasic expression and distorted phase relationships in the epithelium. Log₂ peak-to-trough amplitude and phase of **A)** *Bmal1*, **B)** *Nr1d1*, and **C)** *Dbp* across cell types. Note how epithelial cell types (blue) cluster differently than other tissue cells in both colon regions. Shape size indicates the -log₁₀(p-value) from harmonic regression. Proximal colon (PC): triangles; Distal colon (DC): circles. Log₂ peak-to-trough amplitude and phase for core clock and clock target genes in **D)** Muscularis Mucosa (MM), **E)** Colonocytes_TA, **F)** Enteroendocrine cells. Again the selected epithelial cells show a structure of clock and clock target gene timing compared to MM cells which have a canonical structure. Shape size indicates the -log₁₀(p-value) from harmonic regression. Non-significant genes are indicated by crosses (interpret phase with caution). PC is shown in blue; DC shown in red.

**Supplementary Figure 8.**
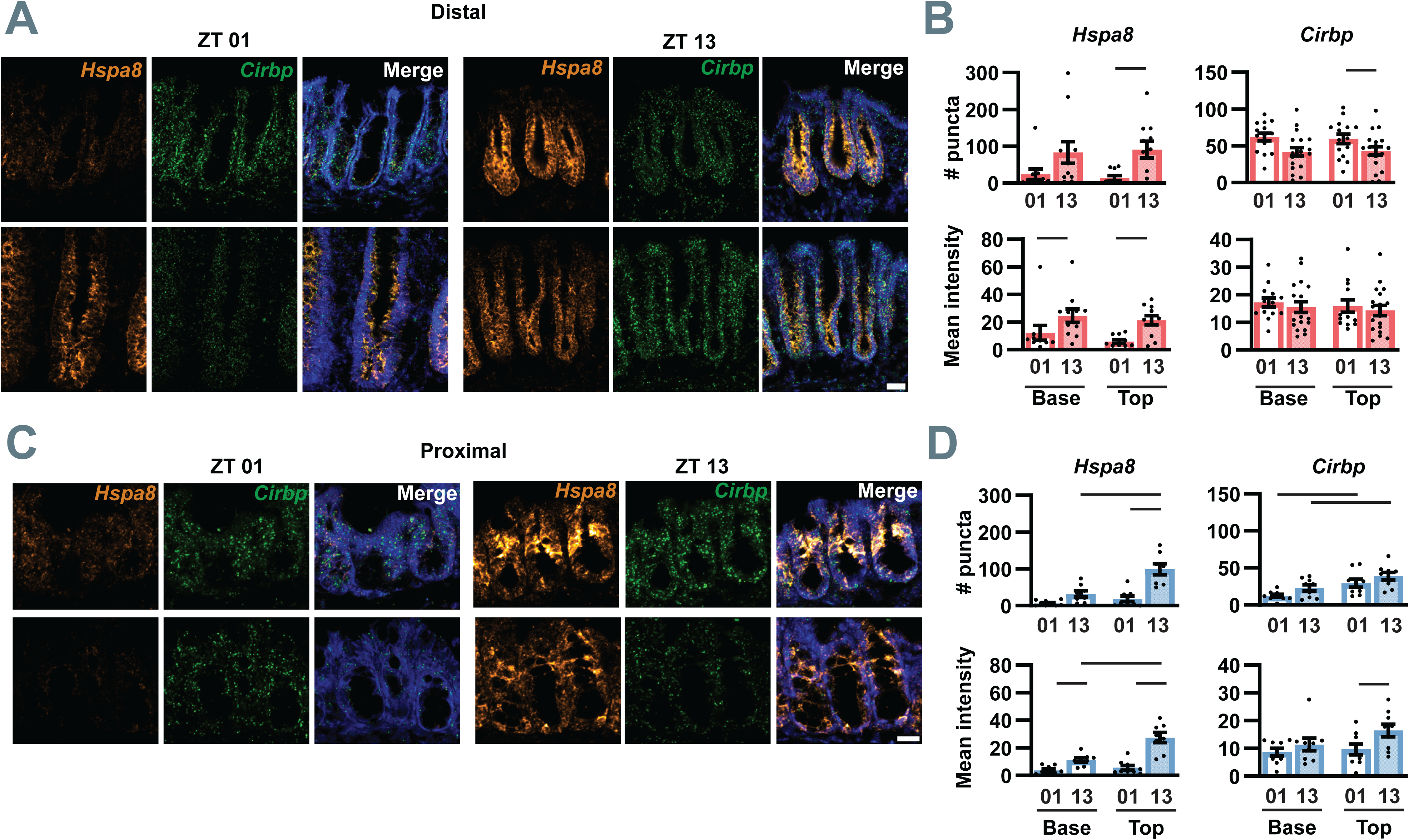
Antiphasic expression of *Hspa8* and *Cirbp* in the distal colon during regeneration. **A-B)** Representative confocal images from *in situ* hybridization showing *Hspa8* and *Cirbp* expression at ZT01 and ZT13 in the proximal colon (scale bar = 20µm). Quantification of *Hspa8* (left) and *Cirbp* (panel) expression in the proximal colon (blue bars) using two methods: counts of RNA puncta at the base and top of the crypt (upper graphs) and mean optical density (lower graphs). C-D) The same imaging and analysis but for the distal colon (scale bar= 25µm) colon **C)** *Hspa8* expression was elevated at ZT13 in both regions while *Cirbp* shows daily variation only in the distal colon, peaking at ZT01. Data are presented as mean ± s.e.m. Significant differences (*p* < 0.05) are indicated by black bars.

## Acknowledgements

We are grateful to technical staff at the Universities of Windsor, Ecole polytechnique fédérale de Lausanne, for help in preparing this study. Genomics was carried out at the Princess Margaret Genomics Centre (https://www.pmgenomics.ca/pmgenomics/).VC-A, JM, ZT, and PK were funded by the Canada Foundation for Innovation, the Ontario Research Fund, and the Canadian Institutes of Health Research. CG, and FN were funded by the Swiss National Science Foundation. To the authors whose research we could not cite due to space limitations, we offer our apologies and thanks.

## Author Contributions

Conceptualization and experimental design was done by VCA, CB, PK, FN; data collection was done by VCA, JM, ZT; data analysis was done by VCA, CB, PK; writing was done by VCA, CB, PK, FN; supervision and project administration was done by PK, FN. All authors have read and agreed to the final version of the manuscript.

